# Red Blood Cells Serve as a Primary Glucose Sink to Improve Glucose Tolerance at Altitude

**DOI:** 10.1101/2025.04.24.650365

**Authors:** Yolanda Marti-Mateos, Ayush D. Midha, Helen Huynh, Will R. Flanigan, Skyler Y. Blume, Isha H. Jain

## Abstract

High altitude conditions result in improved glucose tolerance and lower diabetes risk across species, yet the underlying physiological mechanism remains unclear. Using mouse models, we found that hypoxia alone robustly improved glucose tolerance, independent of insulin sensitivity. This effect persisted for weeks after mice returned to normoxia. PET-CT imaging revealed that internal organs explained only a small fraction of increased glucose uptake in hypoxia, suggesting the presence of an unknown glucose sink. We hypothesized that increased glucose tolerance might be linked to the hypoxia-induced increase in red blood cells (RBCs), whose metabolism relies entirely on glucose. Experimental manipulation of RBC numbers through phlebotomy or transfusion directly altered blood glucose levels, demonstrating the necessity and sufficiency of RBCs as primary glucose sinks in hypoxia. Moreover, RBCs produced during systemic hypoxia exhibited a sustained ∼3-fold increase in glucose uptake, rapidly synthesizing the hemoglobin allosteric regulator 2,3-DPG that allows for increased oxygen release in hypoxia. Therapeutically, we demonstrated that both chronic hypoxia and our recently developed pharmacological hypoxia mimetic, HypoxyStat, effectively rescued hyperglycemia in mouse models of type 1 and type 2 diabetes. Our findings identify RBCs as critical regulators of systemic glucose metabolism under hypoxic conditions, illuminating a conserved physiological adaptation and suggesting novel therapeutic avenues for hyperglycemic disorders.

**Graphical Abstract:** 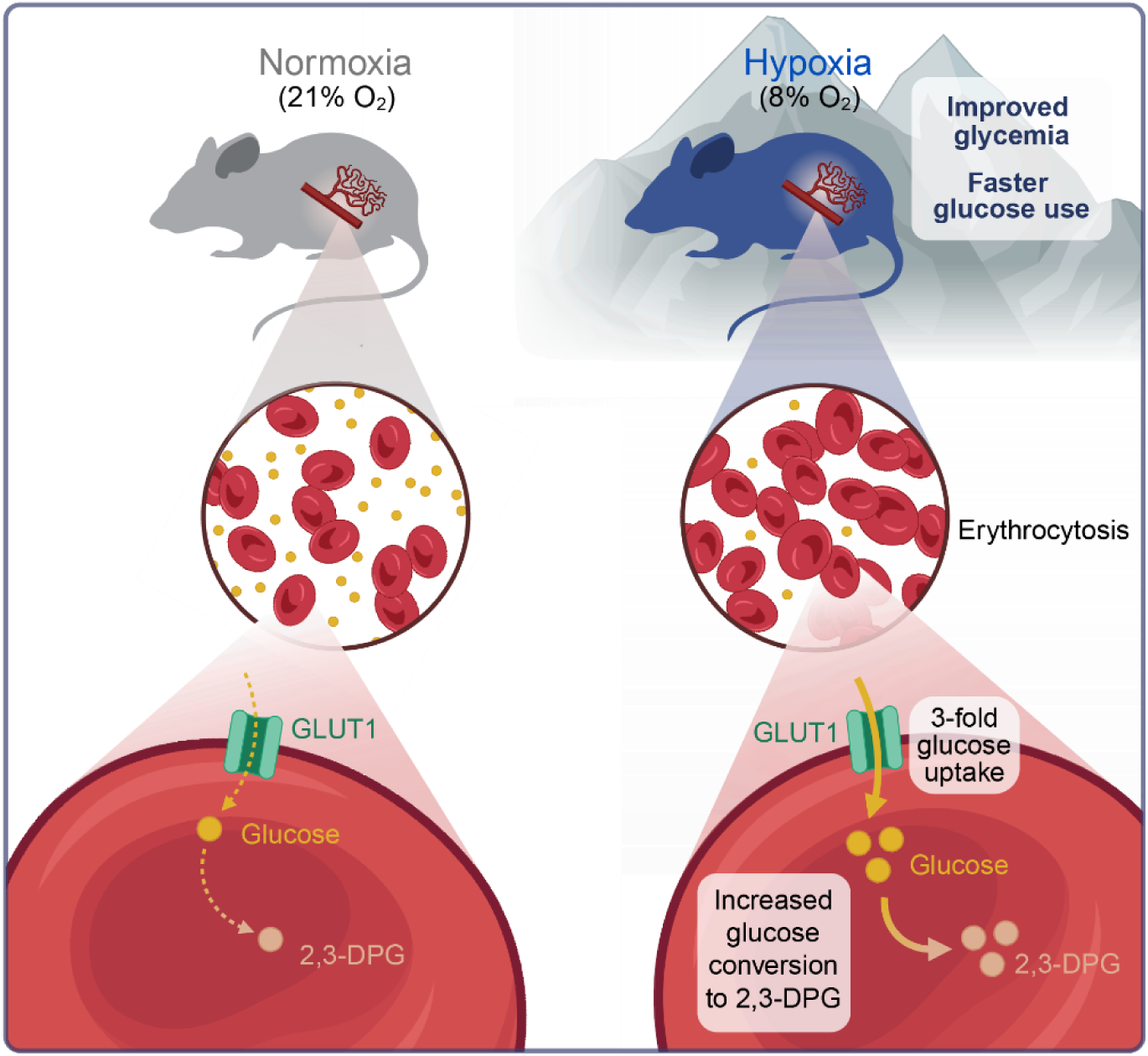

## Introduction

Numerous epidemiological studies report a decreased incidence of diabetes and obesity in high altitude populations^1–7^ (**Table 1**). Improved glucose homeostasis at altitude extends beyond humans. For example, high-altitude-adapted deer mice (*Peromyscus maniculatus*) exhibit enhanced glucose disposal compared to their low-altitude counterparts^8^. Similarly, pigs in Tibet display improved insulin sensitivity and lower plasma glucose concentrations compared to low-altitude breeds^9^, and multiple high-altitude songbirds showed lower glycemia with improved insulin sensitivity^10^. These cross-species observations suggest the existence of an evolutionarily conserved physiological mechanism that optimizes glucose disposal at altitude.

**Table 1.** Human studies linking high altitude with improved glycemic control. Summary of published research articles reporting associations between high-altitude exposure and glucose homeostasis in humans. Included studies report outcomes such as blood glucose levels, glucose tolerance, or diabetes risk, where available. The specific altitude and geographic location of each study are noted.

| Study | Altitude | Location | Glycemia | Glucose Tolerance | Diabetes |
| --- | --- | --- | --- | --- | --- |
| Xu et al., 2017 | 3200-3500m (control)<br>3500m, 4000m | Tibet |  |  | 3500m: -65% risk ↓<br>4000m: -89% risk ↓↓ |
| Woolcott et al., 2015 | 1500-3500m | United States |  |  | 1500-3500: .88 odds ratio ↓ |
| Castillo et al., 2007 | 150m<br>3250m | Peru | 150m: 74.5 mg/dL<br>3250: 50.6 mg/dL ↓ |  |  |
| Calderón et al., 1966 | 4500m | Peru |  | Improved IV glucose tolerance test |  |
| Lindgärde et al., 2004 | 3800m | Peru | Control: 4.6 mM<br>3800m: 3.8 mM ↓ |  |  |
| Lecoultre et al., 2013 | 10 nights at 15% O<sub>2</sub> | Hypoxic tents |  | Increased glucose disposal rate during insulin infusion. |  |
| de Mol et al., 2012 | 12-day trek to 4167m | Morocco | Fasting glucose decreased from 7.6 mM to 6.9 mM ↓ |  | HOMA-IR decreased from 3.8 to 2.7 ↓ |

Many physiological variables change simultaneously at high altitude, making it challenging to determine which factor is specifically responsible for improved glucose homeostasis. High-altitude studies in human populations often lack suitable controls, leaving uncertainty about cause and effect. However, multiple lines of evidence point to decreased oxygen availability (hypoxia) as the critical variable. Some of the earliest systematic investigations into this phenomenon trace back to the pioneering work of the Harvard Fatigue Laboratory between the 1920s and 1940s^11^, which aimed to study human physiology under extreme environmental conditions relevant for soldiers during wartime. In a series of carefully documented experiments, researchers from the Harvard Fatigue Laboratory observed improved glucose tolerance in healthy volunteers who were transported to the Chilean Andes, at elevations reaching up to 6000 meters^11^. These foundational observations have since been revisited and expanded in recent research. For example, our own recent studies demonstrated dramatically reduced blood glucose concentrations in mice exposed chronically to low oxygen, without overt physiological or behavioral deficits.

Despite extensive characterization of improved glucose tolerance at altitude, the precise physiological mechanism underlying this phenomenon remains elusive. A prevailing hypothesis has been that increased peripheral glucose consumption under hypoxic conditions is responsible for this beneficial effect. Indeed, acute hypoxia rapidly upregulates glucose transporters in peripheral tissues via the HIF transcriptional response, thereby enhancing glucose uptake^12,13^. However, these acute signaling responses diminish with prolonged hypoxic exposure due to a negative feedback loop^14^, making it unlikely that such signaling alone can fully explain the sustained improvement in glycemic control observed in chronic hypoxia. Moreover, the gradual onset of improved glycemia over weeks further suggests that slower, chronic adaptations likely contribute to the observed metabolic benefits. Thus, despite numerous studies, the mechanistic basis of long-term improved glycemia at altitude has remained an enduring mystery within the field.

Red blood cells (RBCs) are the most abundant cell type in the human body, accounting for approximately 85% of all cells and contributing to about 4% of total body mass^15^. Under conditions of chronic hypoxia, RBC numbers can nearly double, occupying up to 70% of total blood volume^16^. Mature RBCs are highly specialized cells that lack organelles, such as nuclei and mitochondria. These cells are primarily composed of hemoglobin and play a central role in maintaining adequate tissue oxygenation. Due to their lack of mitochondria, RBC metabolism heavily relies on anaerobic glucose fermentation for ATP production^17^. Interestingly, one of the primary hemoglobin allosteric regulators under hypoxia, 2,3-diphosphoglycerate (2,3-DPG), is continuously produced within RBCs as a glycolytic intermediate via the Rapoport-Luebering shunt^18^. Elevated levels of 2,3-DPG during hypoxia promote enhanced oxygen release from hemoglobin to tissues^19^. These observations suggest an intimate link between RBC glucose metabolism and hypoxic physiological adaptation, raising the intriguing possibility that RBC glucose utilization might play a central role in regulating systemic glucose homeostasis under conditions of low oxygen availability.

Therefore, we hypothesized that RBCs represent the primary glucose sink under hypoxic conditions. To test this hypothesis, we systematically evaluated the relationship between RBC abundance and blood glucose across multiple mouse models in which RBC numbers were experimentally manipulated. Our results revealed that RBC abundance and their altered metabolism in hypoxia could account for most of the observed glycemic changes during hypoxia. These findings uncover an unexpected yet fundamental role for RBCs in systemic glucose homeostasis.

## Results

### Improved Glucose Tolerance Observed in High-Altitude Residents Is Recapitulated in Hypoxic Mice

High-altitude environments are characterized by alterations in several environmental conditions, including lower temperatures, increased UV radiation, decreased humidity, and reduced oxygen availability^20,21^. To determine whether the improved glucose tolerance observed in high-altitude residents results specifically from hypoxia, we employed a mouse model for normobaric hypoxia exposure. We housed eight-week-old male mice under either normoxic (21% O_2_, n=7) or hypoxic (8% O_2_, n=8, corresponding to >5,000 meters of altitude) conditions for three weeks, regularly monitoring blood glucose and body weight. We observed that basal blood glucose levels significantly decreased in hypoxic mice starting from day 2 of exposure, coinciding with acute weight loss that stabilized within one week (**Fig 1A, S1A**). This early reduction in glucose was not explained by reduced food intake, as pair-fed normoxic mice did not show any significant changes in glycemia after three days compared to hypoxic mice (**Fig S1B**). Furthermore, we found glucose tolerance to be substantially improved at all hypoxia timepoints tested (1, 2, and 3 weeks) (**Fig 1B-C**). These results indicate that hypoxia alone recapitulates the enhanced glucose tolerance previously observed in human high-altitude residents.

**Figure 1.**
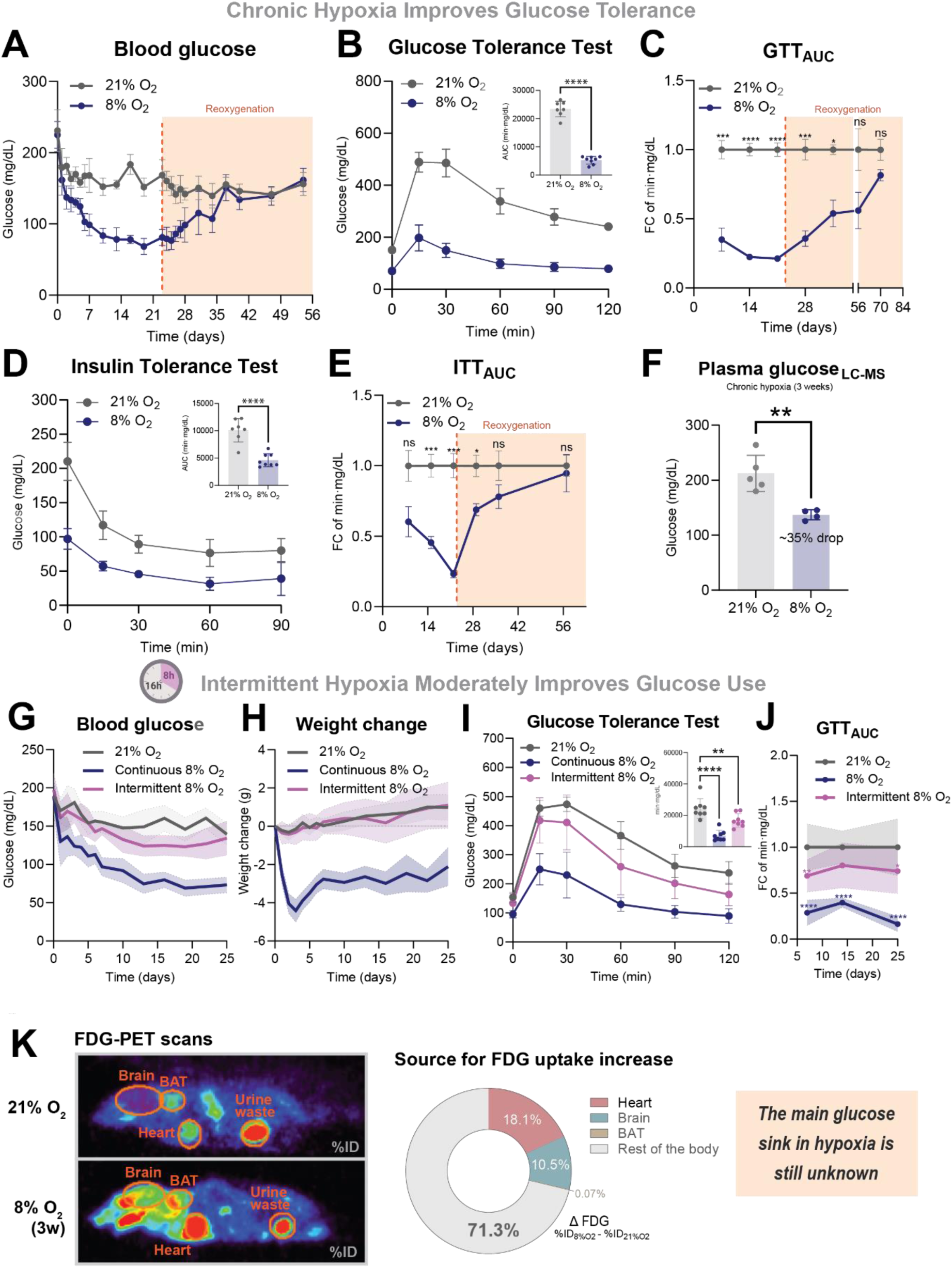
Hypoxia leads to improved glucose tolerance, which is not fully explained by increased glucose uptake by internal organs. **(A)** Tail vein blood glucose levels over time in mice exposed to hypoxia (8% O_2_, n = 8) or normoxia (21% O_2_, n = 7), followed by reoxygenation, indicated by the orange dashed line at day 23. Data are presented as mean ± SD. **(B)** Glucose tolerance test (GTT) performed after two weeks of hypoxia (8% O_2_, n = 8) or normoxia (21% O_2_, n = 7). Data are shown as mean ± SD. Area under the curve (AUC) values from individual GTT curves are plotted (top right) and analyzed using an unpaired two-tailed Student’s t-test. **(C)** Fold change in glucose tolerance, calculated as the AUC from individual tests relative to normoxic controls, in mice under hypoxia (8% O_2_, n = 8) or normoxia (21% O_2_, n = 7), and following reoxygenation. Data are shown as mean ± SEM. Statistical analysis was performed using Sidak’s multiple comparisons test. **(D)** Insulin tolerance test (ITT) after two weeks of hypoxia (8% O_2_, n = 8) or normoxia (21% O_2_, n = 7). Mean ± SD are shown. AUC values from individual ITT curves are plotted (top right) and analyzed using an unpaired two-tailed Student’s t-test. **(E)** Fold change in insulin tolerance, calculated as the AUC from individual tests relative to normoxic controls, in mice exposed to hypoxia (8% O_2_, n = 8) or normoxia (21% O_2_, n = 7), and after reoxygenation. Mean ± SEM are shown. Sidak’s multiple comparisons test was used. **(F)** Plasma glucose concentrations measured by LC-MS in mice exposed to three weeks of hypoxia (8% O_2_, n = 4) or normoxia (21% O_2_, n = 5). Data are presented as mean ± SD. An unpaired two-tailed Student’s t-test was used. **(G)** Tail vein blood glucose levels over time in mice exposed to intermittent hypoxia (8% O_2_ for 8 hours, 21% O_2_ for 16 hours, n = 8), continuous hypoxia (8% O_2_, n = 8), or normoxia (21% O_2_, n = 8). Mean ± SD are shown. **(H)** Body weight changes over time under intermittent hypoxia, continuous hypoxia, or normoxia (all n = 8). Data are shown as mean ± SD. **(I)** Glucose tolerance test after one week of intermittent hypoxia, continuous hypoxia, or normoxia (all n = 8). Mean ± SD are shown. **(J)** Fold change in glucose tolerance over time, quantified as AUC from individual tests relative to normoxic controls, in mice under intermittent hypoxia, continuous hypoxia, or normoxia (n=8 per group). Mean ± SD are shown. Sidak’s multiple comparisons test was used. **(K)** Representative PET-CT scans showing ¹⁸F-FDG signal accumulation in tissues after three weeks of hypoxia (8% O_2_, n = 5) or normoxia (21% O_2_, n = 8). Elliptical ROIs were used to quantify signal in organs including the heart, brain, brown adipose tissue (BAT), and bladder. Bladder signal was subtracted from total body signal to calculate effective whole-body ¹⁸F-FDG accumulation per animal. The difference (Δ%ID₈% - %ID_2_₁%) between individual hypoxic values and the normoxic group average was used to estimate organ-specific contributions to increased glucose uptake. The pie chart displays the relative contributions of individual organs. ∗p < 0.05, ∗∗p < 0.01, ∗∗∗p < 0.001, and ∗∗∗∗p < 0.0001.

Next, to better understand the origin and kinetics of the improved glucose tolerance, we returned mice previously adapted to hypoxia back to normoxic conditions. We regularly monitored blood glucose and body weight to determine when glycemia normalized. Basal blood glucose returned to normoxic levels after 14 days of reoxygenation **(Fig 1A)**. We assessed glucose tolerance after 5, 12, 33, and 47 days of reoxygenation and, surprisingly, found that glucose tolerance did not fully normalize until more than one month after returning to normoxia (**Fig 1B-C, S1C**). These results suggest that the mechanisms responsible for improved glucose tolerance under hypoxic conditions do not require continued hypoxia exposure and persist for weeks after hypoxia cessation.

To determine whether the observed glucose tolerance improvement was insulin-dependent, we performed insulin tolerance tests during both hypoxia adaptation and subsequent reoxygenation. In our model of extreme hypoxia, insulin tolerance did not improve during hypoxia; instead, it gradually decreased over the duration of hypoxic exposure **(Fig 1D-E)**. This phenotype progressively recovered upon reoxygenation, which normalized faster than glucose tolerance **(Fig 1E, S1D)**. Persistent hypoglycemia is known to trigger counter-regulatory mechanisms aimed at preventing dangerously low blood glucose levels^22,23^. As a result, the diminished effect of insulin under these conditions is unlikely to be related to intrinsic insulin resistance at the cellular level. Regardless, these findings support the conclusion that the improvement in glucose tolerance observed in hypoxic mice occurs independently of enhanced insulin signaling.

To determine whether improved glucose tolerance was insulin-dependent, we performed insulin tolerance tests during hypoxia and reoxygenation. Insulin sensitivity declined during hypoxia and gradually recovered with reoxygenation, normalizing more rapidly than glucose tolerance (**Fig. 1D–E, S1D**). As hypoglycemia induces counter-regulatory responses, the reduced insulin effect likely reflects systemic adaptation rather than intrinsic insulin resistance. These findings suggest that glucose tolerance improvements under hypoxia occur independently of enhanced insulin signaling. To assess whether glucose production pathways were contributing to the hypoglycemia phenotype, we conducted a pyruvate tolerance test (**Fig S1E**). Our results indicate that hepatic gluconeogenesis does not significantly contribute to lowered blood glucose during hypoxia, supporting increased glucose clearance as the most plausible explanation.^24–26^

To test whether more subtle hypoxic interventions could similarly improve glucose tolerance, we examined the effects of intermittent hypoxia on glucose homeostasis. We housed eight-week-old WT male mice continuously at 21% O_2_ (continuous normoxia, n=8), continuously at 8% O_2_ (continuous hypoxia, n=8), or intermittently cycling between 21% and 8% O_2_ (intermittent hypoxia, n=8) for 25 days. The intermittent hypoxia protocol consisted of daily cycles with 8 hours of hypoxia exposure during the mice’s sleeping period (9 am to 5pm) and 16 hours of normoxia exposure coinciding with their awake period (5pm to 9am). We regularly monitored blood glucose, body weight, and body temperature. Intermittent hypoxia moderately reduced basal blood glucose (**Fig 1G**) without affecting body weight (**Fig 1H**), indicating that food intake was likely not impaired by sleeping under hypoxic conditions. Consistent with this, body temperature did not acutely decrease under intermittent hypoxia and even showed a tendency to increase during chronic exposure **(Fig S1F)**. Glucose tolerance tests performed at days 7, 14, and 25 revealed a moderate yet consistent improvement throughout the intervention period (**Fig 1I-J**). Taken together, these data suggest that even milder and potentially translational hypoxic interventions, such as intermittent hypoxia therapy, can moderately enhance glucose tolerance.

The above glucose measurements rely on a handheld glucose meter (glucometer). Given the reported technical artifacts of some glucometers when testing blood across wide hematocrit ranges^24–26^, we decided to analyze the plasma fraction alone using LC-MS as an orthogonal method. Plasma glucose levels in mice adapted to hypoxia (8% O_2_) for three weeks showed a 35% decrease compared to normoxic mice, closely matching the 45% reduction previously observed using our glucose meter on whole blood **(Fig 1F, S1G)**. Additionally, we conducted detailed comparisons of glucose concentrations measured by LC-MS (on plasma) and glucose meters (on whole blood or plasma) using matched normoxic and hypoxic samples (**Fig. S1H, S1I**). These analyses revealed a strong correlation between LC-MS and glucose meter quantifications across oxygen tensions. Notably, plasma-based glucose meter readings demonstrated particularly high accuracy (**Fig. S1I**), leading us to adopt this method as our gold standard for glycemia quantification. Collectively, these results validate the accuracy of our glucose measurements and support the reliability of our experimental approach.

### Improved Glucose Tolerance in Hypoxia Is Not Explained by Increased Glucose Uptake by Internal Organs, Suggesting a Missing Glucose Sink

To determine the primary site of glucose clearance under hypoxic conditions, we previously performed PET-CT scans to measure glucose uptake using the tracer 2-deoxy-2-[^18^F]fluoro-D-glucose (FDG)^27^. We injected FDG into mice housed at either normoxia (21% O_2_, n=8) or hypoxia (8% O_2_, n=5) for three weeks. Cellular uptake of radioactive ^18^F-FDG served as a surrogate marker for glucose uptake (**Fig 1K**). Effective whole-body ^18^F signal was quantified in Amide software by placing same-size cylindrical regions of interest (ROIs) around each animal and subtracting FDG accumulation in the bladder, as this fraction represents urinary excretion. To evaluate the contributions of the main organs (brain, heart, BAT) experiencing changes in glucose uptake upon hypoxia exposure^27^, we now quantified ^18^F-FDG accumulation within same-size elliptical ROIs drawn around these organs using CT scans to ensure accurate anatomical identification. We also calculated the difference (Δ_%ID8% - %ID21%_) between whole-body ^18^F signal in hypoxic vs normoxic mice to determine the increase in whole-body glucose uptake due to hypoxia. Surprisingly, we found that internal organs did not account for the majority of hypoxia-driven glucose uptake, with ∼70% of the increase remaining unexplained (**Fig 1K**). These results indicated that the primary glucose sink responsible for enhanced glucose consumption under hypoxic conditions remained unidentified and motivated us to widen our search space.

### Erythrocytosis is Both Necessary and Sufficient to Improve Glycemic Control

Red blood cells (RBCs) are the most abundant cell type in the body, and their total mass increases by approximately 40-50% after four weeks of adaptation to 8% O_2_ exposure **(Fig S2A)**, a phenomenon known as erythrocytosis. Because RBCs lack mitochondria, they primarily metabolize glucose through anaerobic fermentation and the pentose phosphate pathway (PPP)^17^. Previous reports indicate that hypoxic conditions promote glycolytic flux in RBCs over PPP activity^28,29^. Therefore, we hypothesized that the combined effects of increased RBC numbers and enhanced glycolysis in RBCs might account for the majority of the glucose uptake observed under hypoxic conditions.

To directly test if RBCs are the main glucose sink during hypoxia, we reversed erythrocytosis in hypoxic mice via serial phlebotomy and investigated whether this intervention could mitigate the observed decrease in blood glucose (**Fig 2A**). Specifically, we extracted 15% of total blood volume (TBV) every three days from eight-week-old male mice housed under normoxic (21% O_2_, phlebotomized normoxic mice, n=8) or hypoxic (8% O_2_, phlebotomized hypoxic mice, n=8) conditions over four weeks. Hematological and plasma glucose analyses were performed on these mice and compared to control normoxic (n=5) and hypoxic (n=5) mice that had never been subjected to phlebotomy. Phlebotomized hypoxic mice achieved hematocrits comparable to normoxic controls (**Fig 2B, S2B**). Phlebotomy did not significantly alter other blood parameters **(Fig S2C)**. Importantly, this normalization of hematocrit partially normalized the hypoxic hypoglycemia observed in non-bloodlet mice (**Fig 2C-E**), producing a stable glycemic plateau around ∼170 mg/dL. Moreover, glucose tolerance tests performed after four weeks of hypoxia revealed that phlebotomy substantially diminished the improvement in glucose tolerance observed in hypoxic mice (**Fig 2F-G**). Collectively, these data demonstrate that RBC depletion largely normalizes glycemia in hypoxic mice, strongly supporting erythrocytosis as necessary for the enhanced glucose tolerance observed in hypoxia.

**Figure 2.**
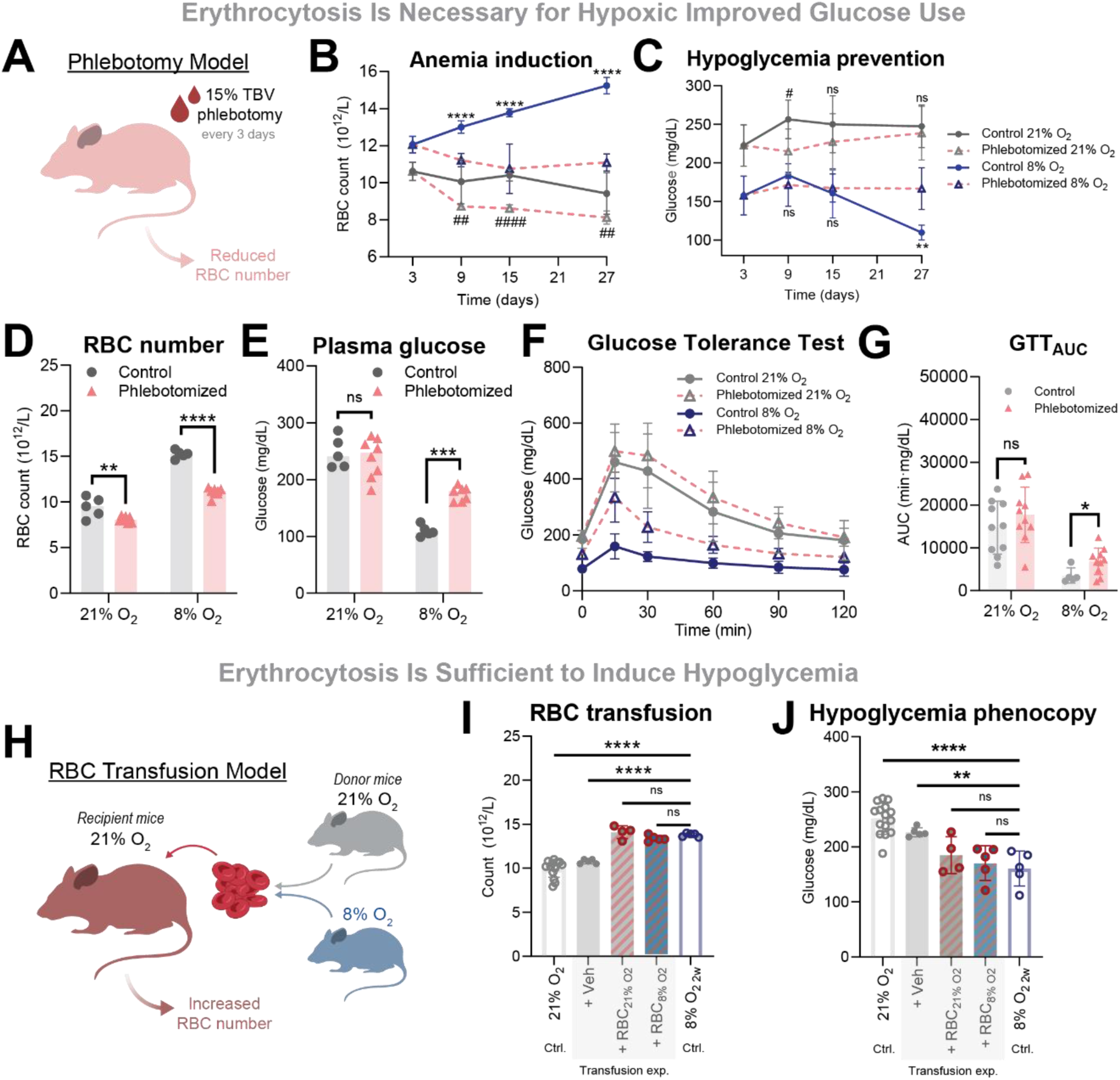
Erythrocytosis is necessary and sufficient to explain hypoglycemia in hypoxia. **(A)** Phlebotomy model. To test whether erythrocytosis is necessary for hypoxia-induced hypoglycemia, red blood cells (RBCs) were depleted by serial phlebotomy in hypoxic (8% O_2_, n = 8) and normoxic (21% O_2_, n = 8) mice. A volume equivalent to 15% of total blood volume (TBV) was removed every three days. Control groups consisted of non-phlebotomized hypoxic (n=5) and normoxic (n=5) mice. **(B)** RBC counts over time in hypoxic (control and phlebotomized, n = 5 and n = 8, respectively) and normoxic (control and phlebotomized, n=5 and n=8, respectively) conditions, demonstrating effective induction of anemia by phlebotomy. Mean ± SD are shown. Two-way ANOVA was used. **(C)** Plasma glucose levels over time in hypoxic and normoxic mice (same groupings as in B), showing the impact of phlebotomy on glycemia. Mean ± SD are shown. Two-way ANOVA was used. **(D)** Endpoint RBC counts after four weeks of hypoxia or normoxia, with or without phlebotomy (n=5–8 per group). Sidak’s multiple comparisons test was used. **(E)** Endpoint plasma glucose levels after four weeks under the same conditions as in D. Sidak’s multiple comparisons test was used. **(F)** Glucose tolerance test (GTT) after four weeks in hypoxic and normoxic mice with or without phlebotomy (n=5–10 per group). Mean ± SD are shown. **(G)** AUC values from individual GTT curves were calculated and analyzed by unpaired two-tailed Student’s t-tests within each oxygen condition. **(H)** RBC transfusion model. To assess whether erythrocytosis is sufficient to induce hypoglycemia, RBCs were isolated from hypoxic (8% O_2_) or normoxic (21% O_2_) donor mice and transfused into normoxic recipients (n=5 per group). An additional control group received vehicle (saline, n = 5). Packed RBCs (75%) were administered twice daily for two consecutive days to rapidly elevate RBC counts. **(I)** Endpoint RBC counts following transfusion with hypoxic, normoxic, or vehicle-derived cells (n=5 per group). For reference, RBC counts from mice exposed to two weeks of hypoxia (8% O_2_) or normoxia (21% O_2_) are included. Ordinary one-way ANOVA was used. **(J)** Endpoint plasma glucose levels after transfusion, with the same grouping and reference data as in I. Ordinary one-way ANOVA was used. ∗p < 0.05, ∗∗p < 0.01, ∗∗∗p < 0.001, and ∗∗∗∗p < 0.0001. (comparisons between phlebotomized and control at 8% O_2_). #p < 0.05, ##p < 0.01, ###p < 0.001, ####p < 0.0001 (comparisons between phlebotomized and control at 21% O_2_).

To directly evaluate whether increased RBC abundance is sufficient to induce hypoglycemia comparable to that seen in hypoxic conditions, we performed RBC transfusions (**Fig 2H**). Specifically, 11-week-old male mice maintained under normoxic conditions (21% O_2_) received two daily retro-orbital injections, for two consecutive days, of either vehicle (saline; n=5) or 75%-packed RBCs derived from donor mice chronically housed at either normoxia (21% O_2_, n=5) or hypoxia (8% O_2_, n=5). Hematological and plasma glucose analyses were conducted one day after the final transfusion round. Transfusion recipients exhibited significant and comparable elevations in RBC count (31% increase with normoxic RBCs, 25% increase with hypoxic RBCs) relative to vehicle-treated controls (**Fig 2I, S2D**). These increments closely matched the RBC elevation observed in mice exposed to hypoxia for two weeks (28% increase). As expected, certain RBC parameters, such as mean corpuscular volume (MCV), differed slightly depending on whether the transfused RBCs originated from normoxic or hypoxic donors, indicating intrinsic differences between RBCs from hypoxic and normoxic mice (**Fig S2D**). Other hematological parameters were not altered (**Fig S2E**). Importantly, plasma glucose levels decreased substantially in normoxic mice transfused with normoxic and hypoxic RBCs (18% and 25% reduction, respectively), closely resembling the 29% glycemic drop observed in mice exposed to two weeks of hypoxia (**Fig 2J**). Together, these findings demonstrate that increased RBC numbers are sufficient to induce hypoglycemia, further substantiating the hypothesis that RBCs represent the primary glucose sink under hypoxic conditions.

### RBCs from Hypoxic Mice Have Increased Glucose Uptake

We next asked whether the observed glucose-lowering effect could be fully explained by the increased RBC number under hypoxia, or if hypoxia additionally enhances glucose uptake per RBC. To address this, we performed experiments using a non-radioactive glucose analog, 2-deoxy-D-glucose (U-^13^C) [2DG (U-^13^C)], as a tracer (**Fig 3A**). Like FDG, 2DG (U-^13^C) is imported into cells through glucose transporters and becomes trapped intracellularly upon phosphorylation, accumulating as 2DG-P (U-^13^C). As this molecule cannot be further metabolized, its accumulation serves as a surrogate marker for glucose uptake, rather than glycolytic flux. We administered 2DG (U-^13^C) retro-orbitally to mice housed under normoxic (21% O_2_, n=5) or hypoxic (8% O_2_, n=5) conditions for three weeks, collecting blood at 2-, 10-, 30-, and 120-minutes post-injection. Plasma 2DG (U-^13^C) concentrations, measured by LC-MS, exhibited similar decay kinetics in both groups, indicating comparable tracer clearance from plasma (**Fig 3B**). However, the accumulation of 2DG-P (U-^13^C) within RBCs occurred much more rapidly in hypoxic mice, reflecting enhanced glucose uptake per RBC (**Fig 3B, S3A**). This phenomenon was even more pronounced when factoring in the increased RBC pool under hypoxic conditions. These results demonstrate that hypoxia increases glucose uptake per RBC *in vivo*, further contributing to the improved glucose tolerance observed during hypoxic exposure.

**Figure 3.**
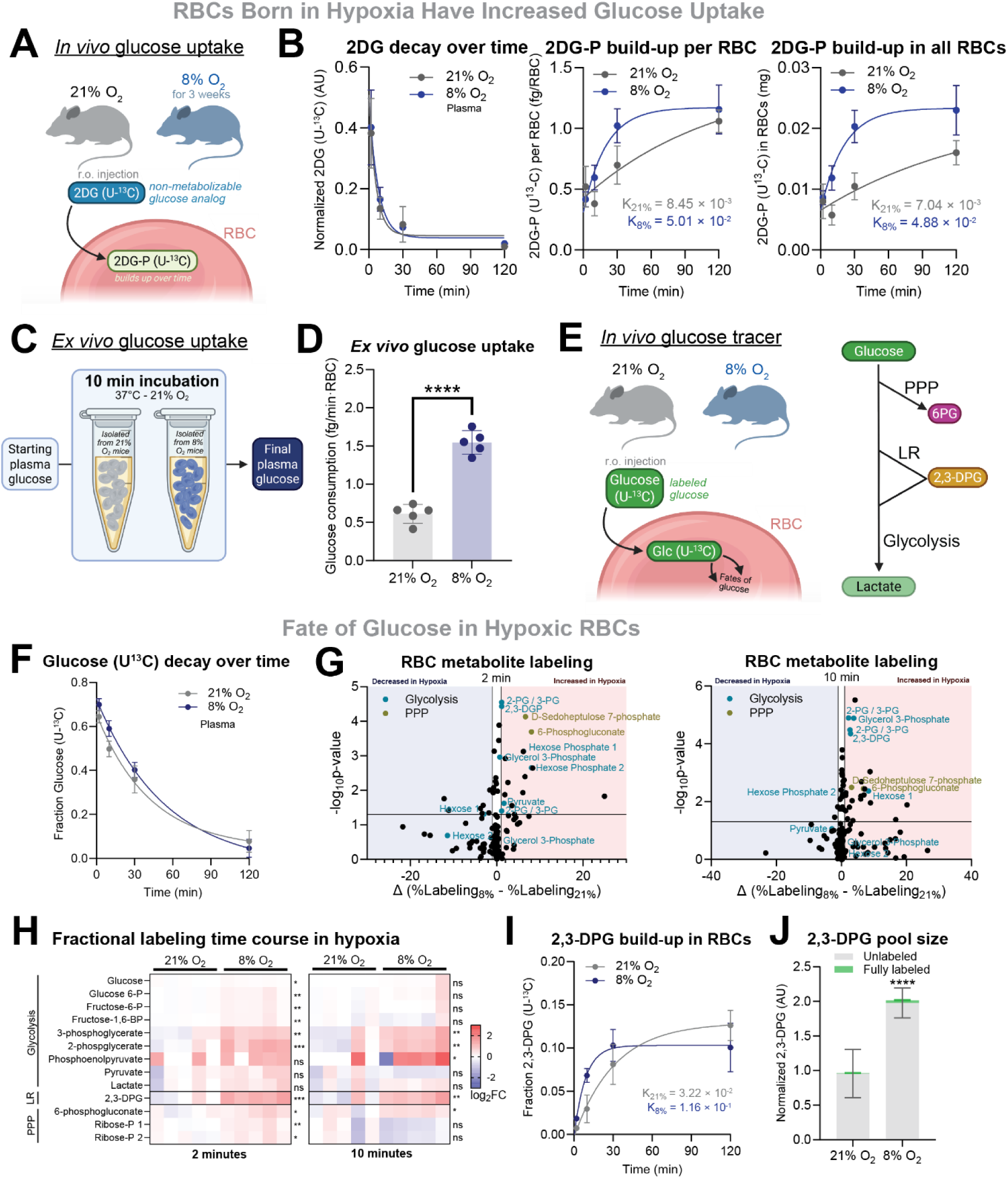
Glucose uptake by RBCs multiplies 3-fold in hypoxia to build hemoglobin allosteric regulator 2,3-DPG. **(A)** *In vivo* glucose uptake experiment. Mice exposed to hypoxia (8% O_2_) or normoxia (21% O_2_) for three weeks received a bolus of uniformly ¹³C-labeled 2-deoxy-D-glucose [2DG (U-¹³C], 1 g/kg body weight via the retro-orbital route. Blood samples were collected at 2-, 10-, 30-, and 120-minutes post-injection. Plasma was analyzed to determine the dynamics of labeled 2-deoxyglucose (2DG), while red blood cells (RBCs) were analyzed for accumulation of phosphorylated 2DG (2DG-P), providing insight into glucose uptake and phosphorylation within RBCs. **(B)** Plasma 2DG decay over time (left), 2DG-P build-up per RBC over time (central) and 2DG-P estimated accumulation in all the RBC pool over time (right) in mice exposed to hypoxia (8% O_2_, n=5) or normoxia (21% O_2_, n=5) for three weeks. Normalization was done by dividing the desired peak area by an internal unlabeled 2DG-P standard, giving arbitrary units (AU) for 2DG but precise concentrations for 2DG-P measurements. 2DG-P quantifications were further normalized by analyzed RBC number per sample (**Fig S3A**) and estimated total RBC number per animal assuming an equal total blood volume of 1.5 mL across mice. **(C)** *Ex vivo* glucose uptake experiment. Blood was collected from mice exposed to hypoxia (8% O_2_) or normoxia (21% O_2_) for four weeks. RBCs were isolated and incubated with pooled normoxic plasma of a known glucose concentration for 10 minutes at 37°C. Glucose uptake was quantified by measuring the drop in plasma glucose levels post-incubation, normalized to RBC number per sample (**Fig S3B**). A plasma-only control was included to confirm that glucose reduction was RBC-dependent. **(D)** *Ex vivo* glucose uptake per RBC. Assayed RBCs were isolated from mice exposed to hypoxia (8% O_2_, n=5) or normoxia (21% O_2_, n=5) for four weeks. **(E)** Glucose tracer experiment. Mice exposed to hypoxia (8% O_2_) or normoxia (21% O_2_) for three weeks received a bolus of uniformly labeled glucose (U-¹³C) (1 g/kg). Blood was collected at multiple time points post-injection, and plasma and RBCs were isolated and stored at −80°C for downstream isotopic analysis. **(F)** Plasma fractional glucose labeling over time post-injection in mice exposed to hypoxia (8% O_2_, n=5) or normoxia (21% O_2_, n=5) for three weeks. **(G)** Volcano plots depicting the delta in fractional labeling in hypoxia to normoxia (Δ (%Labeling8% - %Labeling21%) and their p-value after unpaired Student’s t-test analysis from the semi-targeted metabolomics analysis on glucose-derived metabolites at 2- (left) and 10-minutes (right) post-injection. Glycolytic (blue) and PPP (brown) intermediates are highlighted in the graph. **(H)** Heatmap from targeted metabolomics analysis integrating the main glucose-derived metabolites from glycolysis, LR and PPP in RBCs. Statistical significance of log2FC are depicted on the right of the heatmap. **(I)** 2,3-DPG fractional labeling in RBCs over time post-injection in mice exposed to hypoxia (8% O_2_, n=5) or normoxia (21% O_2_, n=5) for three weeks. **(J)** Unlabeled and fully labeled 2,3-DPG at the earliest timepoint, 2 minutes after the tracer injection, as a qualitative indication for 2-DPG pool size. Normalization was done by dividing the peak area of these metabolites to Val-d8 peak area (AU). Uncorrected Fisher’s LSD multiple comparisons was used. ∗p < 0.05, ∗∗p < 0.01, ∗∗∗p < 0.001, and ∗∗∗∗p < 0.0001.

As an orthogonal approach, we further evaluated RBC glucose uptake using an *ex vivo* system. Blood was collected from mice exposed to normoxia (21% O_2_, n=5) or hypoxia (8% O_2_, n=5) for four weeks, and RBCs were isolated by centrifugation. Identical volumes of RBCs from normoxic or hypoxic mice were incubated *ex vivo* with equal volumes of plasma (1:1 RBC-to-plasma ratio), standardized to an identical initial glucose concentration (**Fig 3C**). After 10 minutes of incubation at 37°C under normoxic conditions, we measured residual glucose in the plasma to determine glucose uptake. Hematology analyses were performed concurrently to accurately quantify the RBC number in each sample (**Fig S3B**). RBCs derived from hypoxic mice exhibited a 2.5-fold increase in glucose uptake per cell compared to normoxic controls (**Fig 3D**). Importantly, because this experiment was conducted entirely under normoxic conditions, we conclude that the enhanced glucose uptake observed in hypoxia-derived RBCs reflects a sustained functional adaptation rather than a hyperacute response to low oxygen availability. Thus, our *ex vivo* data further substantiates that chronic hypoxia induces a long-lasting increase in glucose uptake capacity per individual RBC.

Having confirmed that glucose uptake is markedly increased in RBCs from hypoxic mice and that glucose synthetic pathways are likely not affected in hypoxia (**Fig S1E**), we next sought to determine the metabolic fate of this enhanced glucose influx. To address this, we performed an *in vivo* tracing experiment using uniformly labeled D-glucose (U-^13^C). We injected glucose (U-^13^C) retro-orbitally into mice exposed to normoxia (21% O_2_, n=5) or hypoxia (8% O_2_, n=5) for three weeks and collected blood at 2-, 10-, 30-, and 120-minutes post-injection (**Fig 3E**). Plasma glucose (U-^13^C) levels over time and RBC metabolites were analyzed by LC-MS. As expected, labeled glucose in plasma decayed promptly over time (**Fig 3F**), implying rapid disposal by cells. In order to profile glucose fates in RBCs under hypoxic conditions comprehensively, we conducted semi-targeted metabolomics (**Fig 3G**). This analysis revealed that 2,3-DPG—the primary product of the LR pathway—and glycolytic intermediates were the metabolites whose labeling was most significantly increased in RBCs from hypoxic mice (**Fig 3G**). Some other metabolites whose labeling from glucose increased in hypoxia were gluconate and α-D-Glucose 1,6-bisphosphate, while others like phosphonoacetate were decreased.

Targeted quantification of glycolytic, Luebering-Rapoport (LR), and pentose phosphate pathway (PPP) intermediates confirmed a marked accumulation of labeled 2,3-DPG, along with other glycolytic intermediates (**Fig. 3H**). Importantly, 2,3-DPG serves as a key allosteric regulator of hemoglobin, facilitating enhanced oxygen release to hypoxic tissues ^19^. Fractional labeling analyses revealed a ∼3.5-fold increase in the rate constant of 2,3-DPG in hypoxic versus normoxic RBCs (**Fig. 3I**). While precise flux calculations require 2,3-DPG pool size quantification, the observed 2-fold increase in unlabeled and 5-fold increase in fully labeled 2,3-DPG under hypoxia (**Fig. 3J**) indicate a qualitatively higher metabolic flux. In summary, our data demonstrate that hypoxia markedly enhances *in vivo* glucose utilization by RBCs, directing it primarily toward the synthesis of 2,3-DPG, a hemoglobin modulator, as well as other glycolytic intermediates.

### Hypoxia Rescues Hyperglycemia in Type 1 Diabetes Model

To investigate whether the hypoxia-induced increase in RBC number and metabolic rewiring could serve as a therapeutic approach for hyperglycemia, we tested the efficacy of hypoxia in treating elevated blood glucose using two different disease models. First, we evaluated hypoxia’s potential as a therapy for hyperglycemia induced by type I diabetes (**Fig 4A**). We injected 8-week-old male mice intraperitoneally with either vehicle or streptozotocin (STZ), a compound that leads to loss of insulin-producing pancreatic β-cells^30^, causing insulin deficiency and hyperglycemia (**Fig S4A**). To better replicate the clinical timeline, hypoxia therapy was initiated in diabetic mice only after the diagnosis of diabetes was confirmed, two weeks after the final STZ administration. STZ- and vehicle-treated mice were then randomized into normoxic (21% O_2_; vehicle-treated, n=8; STZ-treated, n=8) or hypoxic therapy (8% O_2_; vehicle-treated, n=8; STZ-treated, n=8) conditions for three weeks. Hypoxic exposure induced erythrocytosis in both STZ-treated and vehicle-treated mice (**Fig 4C, S4D**), whereas RBC counts remained unchanged under normoxic conditions. Basal blood glucose levels and body weight were monitored every three days throughout the study period. Notably, hypoxic STZ-treated mice exhibited a dramatic reduction in their initially elevated glycemia (**Fig 4B, S4B-C)**. Glucose tolerance tests performed after three weeks demonstrated that hypoxia completely rescued impaired glucose tolerance caused by STZ treatment, highlighting the therapeutic potential of our intervention (**Fig 4D-E**).

**Figure 4.**
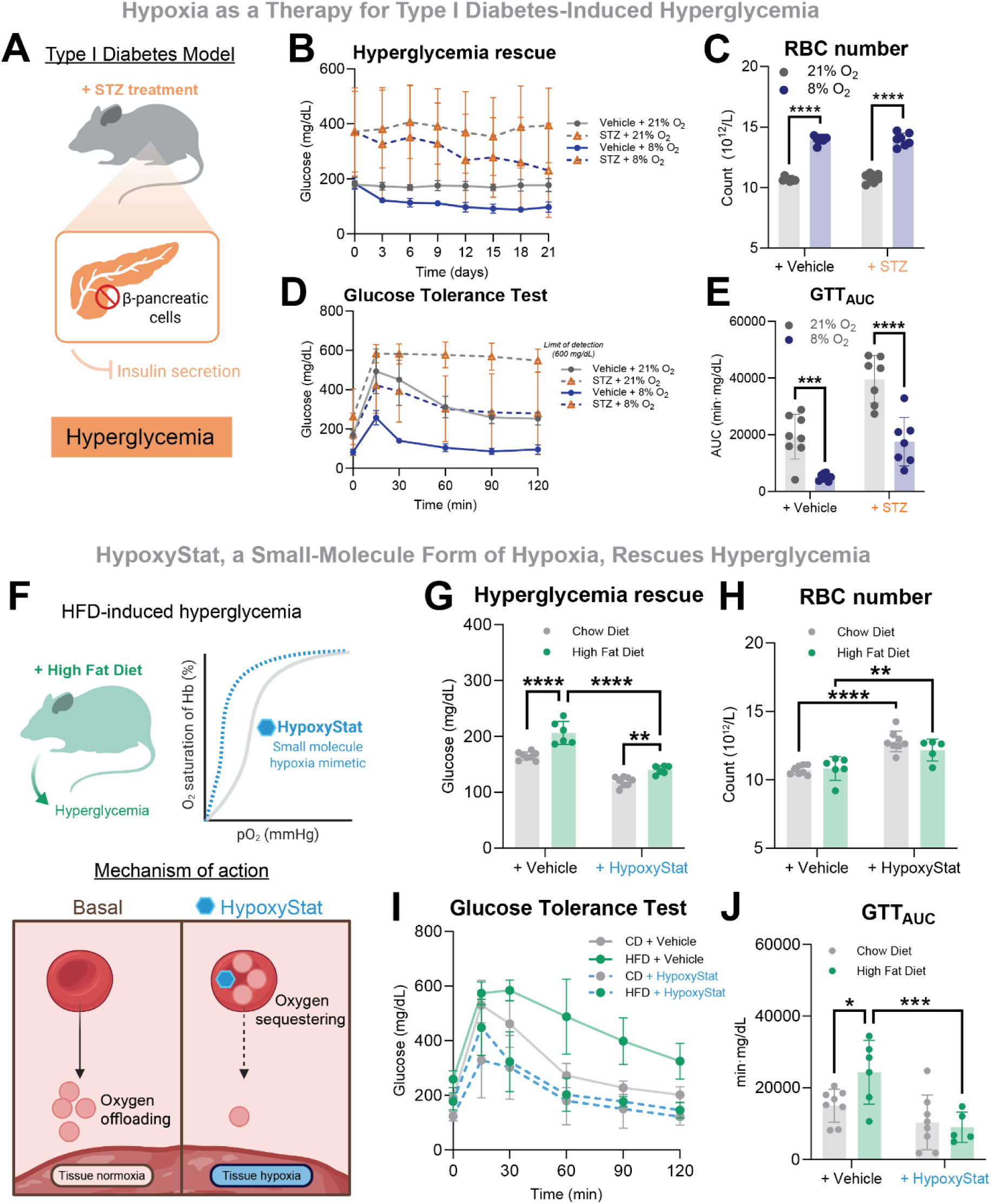
Hypoxia ameliorates STZ- and HFD-induced hyperglycemia. **(A)** Type I Diabetes Model. Type I diabetes was induced in male mice via five consecutive daily i.p. injections of streptozotocin (STZ) or vehicle. Two weeks post-injection, hyperglycemia was confirmed, and mice were randomized to hypoxia (8% O_2_) or normoxia (21% O_2_) for three weeks. Blood glucose and body weight were monitored throughout to assess the impact of hypoxia in diabetic and control mice. **(B)** Tail vein blood glucose over time in hypoxia (8% O_2_ Vehicle- or STZ-treated mice, n=8 and n=8, respectively) or normoxia (21% O_2_ Vehicle- or STZ-treated mice, n=8 and n=8, respectively). Mean ± SD are shown. **(C)** Endpoint RBC count after three weeks in hypoxia (8% O_2_ Vehicle- or STZ-treated mice, n=8 and n=7, respectively) or normoxia (21% O_2_ Vehicle- or STZ-treated mice, n=8 and n=8, respectively). Sidak’s multiple comparison test was used. **(D)** Glucose tolerance test after three weeks in hypoxia (8% O_2_ Vehicle- or STZ-treated mice, n=8 and n=7, respectively) or normoxia (21% O_2_ Vehicle- or STZ-treated mice, n=8 and n=8, respectively). Mean ± SD are shown. **(E)** AUC values from individual GTT curves were plotted and analyzed by Sidak’s multiple comparison test. **(F)** Model for HFD-Induced Hyperglycemia. Male C57BL/6J mice fed either a high-fat diet (HFD) or standard chow diet (CD) were treated daily with HypoxyStat (600 mg/kg) or vehicle for 2.5 weeks. Briefly, HypoxyStat increases hemoglobin’s oxygen affinity, limiting oxygen release and inducing tissue hypoxia. HFD-fed mice received HypoxyStat (n=5) or vehicle (n=6), and CD-fed mice received HypoxyStat (n=8) or vehicle (n=8). Blood glucose was measured after two weeks, and glucose tolerance was assessed after 2.5 weeks to evaluate the metabolic effects of HypoxyStat under different dietary conditions. **(G)** Tail vein blood glucose in HypoxyStat-treated mice (HypoxyStat CD- or HFD-fed, n=8 and n=5, respectively) or vehicle-treated animals (Vehicle CD- or HFD-fed, n=8 and n=6, respectively). Sidak’s multiple comparison test was used. **(H)** Endpoint RBC count in HypoxyStat-treated mice (HypoxyStat CD- or HFD-fed, n=8 and n=5, respectively) or vehicle-treated animals (Vehicle CD- or HFD-fed, n=8 and n=6, respectively). Sidak’s multiple comparison test was used. **(I)** Glucose tolerance test after HypoxyStat dosing (HypoxyStat with CD or HFD, n=8 and n=5, respectively) or vehicle-dosing (Vehicle with CD or HFD, n=8 and n=6, respectively). Mean ± SD are shown. **(J)** AUC values from individual GTT curves were plotted and analyzed by Sidak’s multiple comparison test. ∗p < 0.05, ∗∗p < 0.01, ∗∗∗p < 0.001, and ∗∗∗∗p < 0.0001.

Second, we evaluated whether HypoxyStat, a small-molecule compound we recently developed to pharmacologically cause hypoxia^31^, could effectively reverse hyperglycemia induced by high-fat diet (HFD) (**Fig 4F**). Briefly, HypoxyStat increases hemoglobin’s oxygen affinity, limiting oxygen offloading to the tissues and inducing local tissue hypoxia^31^. Male mice fed either chow diet (CD) or HFD received daily doses of vehicle or HypoxyStat via oral gavage, all while being housed under normoxic conditions (21% O_2_). As expected, vehicle-treated HFD-fed mice developed prominent hyperglycemia compared to CD-fed controls (**Fig 4G, S4E**). Notably, HypoxyStat treatment completely abolished HFD-induced hyperglycemia (**Fig 4G, S4E**), coinciding with a robust induction of erythrocytosis (**Fig 4H, S4F**). Importantly, no other blood parameters were affected (**Fig S4G**). Glucose tolerance was significantly impaired in vehicle-treated HFD-fed mice but fully normalized upon treatment with HypoxyStat, irrespective of dietary fat content (**Fig 4I-J**). Collectively, these findings demonstrate that HypoxyStat, our recently developed pharmacological hypoxia mimetic, potently rescues hyperglycemia and impaired glucose tolerance associated with diet-induced obesity.

## Discussion

Reduced glycemia, improved glucose tolerance, and lower risk for diabetes and obesity have long been observed among high-altitude populations^1–7^ and across diverse organisms^8–10^, yet the physiological mechanisms underlying these beneficial effects on glucose metabolism have remained unclear. Here, we identify red blood cells (RBCs) as the principal glucose sink during hypoxia (**Fig 5**). Hypoxia triggers erythrocytosis, a physiological increase in RBC number of up to ∼50%, and these newly generated RBCs display a markedly enhanced capacity for glucose uptake and metabolism. Indeed, we demonstrate that depletion of RBCs in hypoxic mice effectively normalizes blood glucose, while artificially increasing RBC number under normoxic conditions is sufficient to induce hypoglycemia. At the cellular level, hypoxia boosts glucose uptake per individual RBC approximately 3-fold, enabling rapid accumulation of the hemoglobin allosteric regulator 2,3-diphosphoglycerate (2,3-DPG), a critical molecule for oxygen offloading and hypoxia adaptation^19^. These findings highlight a previously unrecognized physiological mechanism and could inspire innovative therapeutic strategies for treating hyperglycemia, as evidenced by the efficacy of our hypoxia-based interventions.

**Figure 5.**
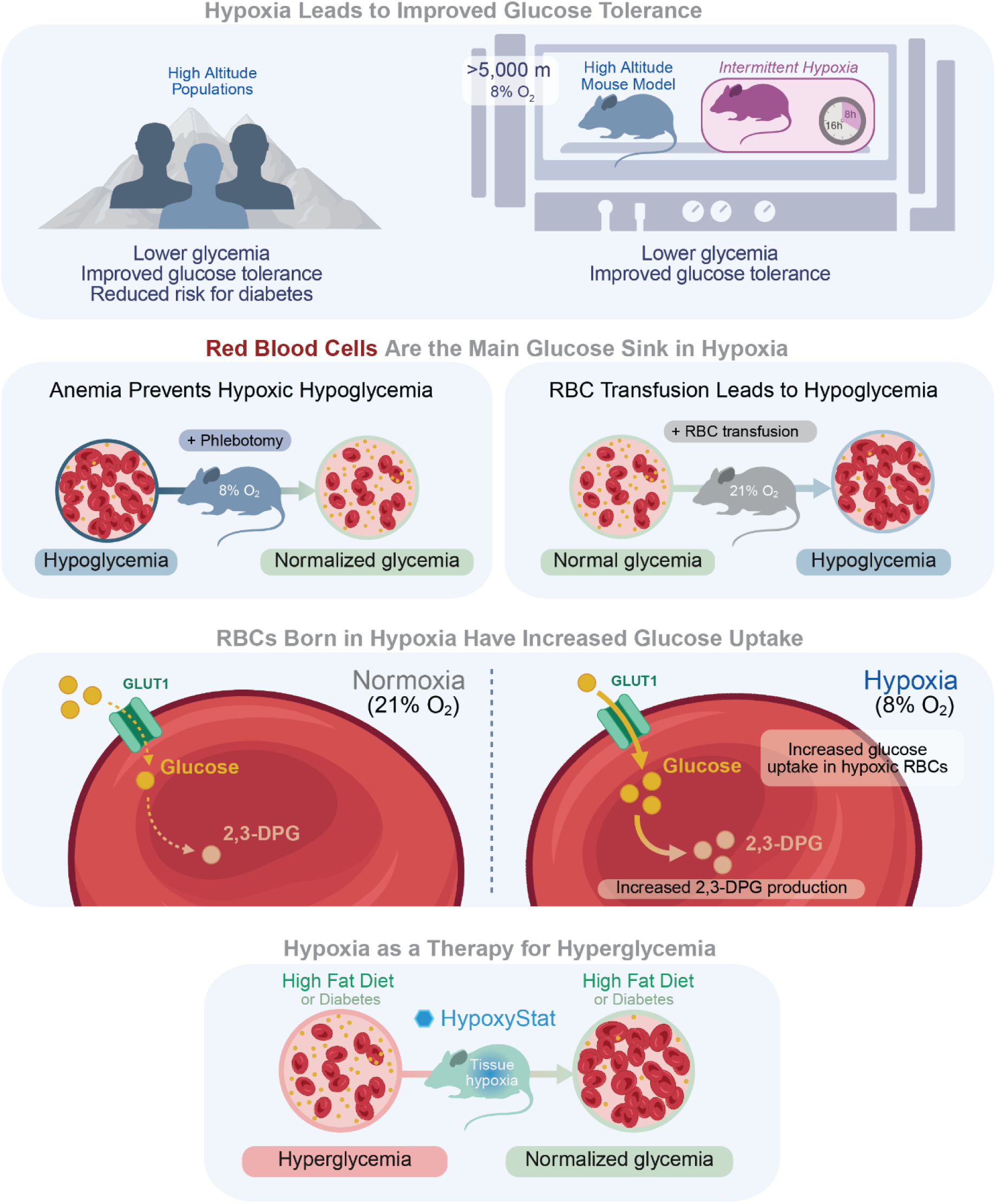
Summary of the physiological mechanism behind hypoxic hypoglycemia. Hypoxia improves glucose tolerance at high altitude, but the mechanism has remained unclear. In mice, we show that hypoxia alone lowers glycemia and enhances glucose tolerance. To assess whether RBCs are a key glucose sink in hypoxia, we manipulated RBC numbers which in turn altered blood glucose levels. We also showed that RBCs produced under hypoxia presented a sustained increase in glucose uptake and 2,3-DPG production. Importantly, pharmacological hypoxia mimetic (HypoxyStat) reversed hyperglycemia in diabetic mouse models. Altogether, these findings identify RBCs as central players in hypoxia-driven glucose regulation and highlight a potential therapeutic pathway for diabetes.

The disproportionately large effect of RBC abundance on glucose homeostasis in hypoxia may partly be explained by a shift toward a younger, metabolically-active RBC population. In our phlebotomy model, we prevented approximately 80% of the hypoxia-induced RBC increase, yet this intervention reduced the glycemia drop by about 55%. This apparent discrepancy can likely be explained by considering RBC population dynamics. Under normal conditions, erythropoiesis occurs at a minimal rate^32^, resulting in a predominantly aged RBC population with only a small fraction of newly synthesized, younger cells. Younger RBCs, however, are functionally distinct, exhibiting significantly higher glucose use than older RBCs^33^. Importantly, both hypoxia and phlebotomy strongly stimulate erythropoiesis^34^, thus shifting the RBC population towards younger, glucose-avid cells. Consequently, the 50% hematocrit observed in phlebotomized hypoxic mice is not directly comparable to the ∼45% hematocrit found in normoxic controls, because the former group contains a much younger RBC population with enhanced glucose metabolism.

The enhanced glucose uptake capacity of younger RBCs observed under hypoxia may be explained by increased GLUT1 expression established during erythropoiesis. GLUT1 is the primary glucose transporter in mature RBCs^35,36^, although some studies have indicated minimal expression of alternative glucose transporters, such as GLUT4^37^ and GLUT3^38^. Given that mature RBCs are enucleated and thus unable to synthesize new proteins, their glucose uptake capacity largely reflects passive factors established at earlier developmental stages. These include GLUT1 expression levels determined at the progenitor cell stage^39^, as well as structural modifications influencing membrane fluidity, the redox state of specific residues, interactions with other proteins, or binding of allosteric regulators^40,41^. GLUT1 expression in erythroid progenitors is known to be regulated by the hypoxia-inducible factor (HIF) pathway^42–44^, which becomes activated during erythropoiesis and under hypoxic conditions. Because the half-life of GLUT1 protein in RBCs is thought to approach the RBC lifespan itself, increased glucose uptake observed under hypoxia may be largely attributable to an expanded pool of younger RBCs, which retain higher GLUT1 expression established during their recent erythropoietic development^39^. Future work will dissect the contribution of RBC age and GLUT1 expression/localization to the improved glycemia during systemic hypoxia.

Beyond enhanced glucose uptake, we observed a marked acceleration in the conversion of glucose to 2,3-DPG and other glycolytic intermediates during hypoxia. These findings align with the recently reported role of the Band 3 metabolon in regulating the compartmentalization of glycolytic enzymes in response to oxygen availability^45,46^. Under normoxic conditions, Band 3 (an RBC plasma membrane protein) sequesters glycolytic enzymes, thereby promoting flux through the PPP to support antioxidant defense mechanisms. In contrast, under hypoxia, deoxygenated hemoglobin displaces these enzymes from Band 3, thereby enhancing glycolytic flux and favoring the accumulation of 2,3-DPG. This shift is consistent with our metabolic profiling and supports a functional adaptation of red blood cell metabolism to hypoxia.

Our findings strongly support the idea that hypoxic hypoglycemia primarily results from accelerated and enhanced glucose uptake by red blood cells. Nevertheless, changes in gluconeogenesis, glycogenolysis, or glucose absorption and secretion might also contribute to systemic hypoglycemia. Preliminary data from pyruvate tolerance tests, which serve as a proxy for hepatic gluconeogenic capacity, indicated no significant differences between oxygen conditions. However, further investigation into additional pathways, including glycogen breakdown and intestinal glucose absorption, is warranted. This will enable a comprehensive understanding of hypoxia’s influence on glucose metabolism beyond its clearly demonstrated effects on glucose uptake.

Glucose uptake per individual RBC was increased in hypoxia, but the mechanism did not depend on acute low oxygen exposure, as the difference was still observed under constant room air in our *ex vivo* experimental system. This finding is consistent with our data on glycemia recovery dynamics upon reoxygenation, as the beneficial effects of hypoxia were long maintained for the first month after re-exposure to normoxic air. Murine RBC half-life ranges from 16.5 to 30 days^47^. Thus, the glycemia normalization upon reoxygenation could be explained by the degradation and turnover of the newly synthesized RBCs. Of note, this phenotype lies in contrast to our previous work on developing hypoxia as a therapy for mitochondrial diseases^31,48–50^. In that setting, chronic and continuous hypoxia was required for sustained benefit and return to normoxic conditions led to rapid decline of the premiere mitochondrial disease mouse model. By contrast, the beneficial effects of hypoxia on glycemia appear to be sustained for weeks after exposure which bodes well for translational approaches.

Our data also offer insights into broader population-level observations, clarifying previously unexplained differences in glycemic control among high-altitude residents^1–7^. Most high-altitude populations consistently demonstrate improved glucose tolerance; however, Sherpas represent a notable exception, as they do not exhibit improved glycemic control despite chronic high-altitude exposure^16,51,52^. Genetically, Sherpas carry hypoxia-adaptive variants in HIF2 (*EPAS1*) and related genes that blunt hypoxia-induced erythropoiesis, resulting in unusually low hematocrit levels compared to other high-altitude groups. Similarly, among Andean populations, individuals carrying the *EPAS1*/*HIF2* missense variant rs570553380 also exhibit reduced hematocrit^53,54^. Our results provide a clear mechanistic explanation for this discrepancy: the lack of an expanded pool of newly synthesized, glucose-avid RBCs likely prevents Sherpas from experiencing the glycemic benefits typical of other highlanders.

Our findings are also relevant beyond high altitude residents. In fact, it has been reported that Chuvash polycythemia patients, who present marked erythrocytosis due to sustained HIF signaling in basal conditions, have decreased blood glucose levels, in agreement with our work. In a similar context, erythropoietin (EPO) treatment has been associated with improved glucose tolerance and reduced blood glucose levels in both human subjects^55^ and animal models^56–58^. Conversely, studies have shown that anemic patients often present with hyperglycemia^59^ and that diabetes is frequently associated with anemia^60–64^, further supporting our proposed physiological mechanism. Collectively, these scattered yet converging observations suggest that the role of red blood cells in glycemic control has been largely overlooked and may carry broader clinical implications.

A future therapeutic avenue against hyperglycemia could be the use of our small-molecule form of hypoxia, HypoxyStat^31^. However, the risks associated with increased blood viscosity may counter the benefits for a relatively manageable condition such as diabetes. A related approach would entail promoting a faster turnover of RBCs that shifted RBC pools towards younger, glucose-avid RBCs. Such an intervention would not affect blood viscosity while potentially improving glycemic control.

In summary, we have identified red blood cells as key regulators of systemic glucose metabolism under hypoxic conditions. Of note, related findings were recently reported by Scherer et al.^65^ further corroborating our observations. Increased RBC abundance and metabolic reprogramming dramatically enhance glucose uptake, positioning RBCs as mobile glucose reservoirs. This discovery clarifies longstanding observations of improved glycemic control among high-altitude residents and explains clinical links between RBC number and blood glucose in polycythemia and anemia. Our findings thus open new therapeutic avenues, including the modulation of RBC dynamics or use of pharmacological hypoxia mimetics such as HypoxyStat, for effectively treating hyperglycemia and associated metabolic disorders.

## Materials & Methods

### Animal model

Male C57BL/6J (#000664) mice (8-11 weeks old) from The Jackson Laboratory were used for all animal experiments. Mice were housed at 24°C on a 12h:12h light:dark cycle. Unless stated differently, mice were fed a chow diet (PicoLab 5058). Cages were randomly allocated to normoxic or hypoxic chambers within the Gladstone Institutes animal facility. Hypoxia was simulated in chambers by mixing N_2_ (Airgas), O_2_ (Airgas, Praxair), and room air using gas regulators. FiO_2_ and CO_2_ levels were continuously monitored wirelessly and checked daily. To inhibit the accumulation of CO_2_, soda lime (Fisher Scientific) was placed surrounding cages in each chamber. Mouse experiments were approved by the UCSF Institutional Animal Care & Use Program (IACUC).

### Body weight measurements

Body weight was monitored daily during the first week of hypoxia exposure and every three days during the following weeks of treatment, unless specified differently.

### Pair-feeding experiment

Food intake per mouse was calculated by measuring the daily change in food weight in single-housed cages. For the pair-fed group, food availability was restricted to match the amount consumed by hypoxic mice on the previous day, resulting in a one-day temporal lag between the hypoxic (8% O_2_, n=8) group and the normoxic (21% O_2_) pair-fed and ad libitum-fed (n=8 and n=8, respectively) groups. All mice were single-housed for the whole experiment. Body weight was recorded daily and blood glucose measurements were performed on whole blood using the OneTouch Ultra Plus^TM^ glucose meter.

### Blood and plasma glucose measurements

Blood glucose levels were measured in mice during their sleeping phase (9:00 AM to 12:00 PM) without prior fasting interventions. Given that mice predominantly feed during their active (dark) phase (6:00 PM to 6:00 AM), we presume that these measurements were obtained after at least three hours of physiological fasting. In experiments involving multiple longitudinal blood glucose measurements, the time of sampling was kept consistent across all time points.

For standard blood glucose analysis, a small drop of whole blood was obtained via a gentle tail tip incision and immediately measured using the OneTouch Ultra Plus™ glucose meter. Although we also validated the OneTouch Verio™ glucose meter, all reported experiments utilized the Ultra Plus™ version.

To account for potential measurement inaccuracies due to elevated hematocrit levels, we confirmed that applying the plasma fraction directly to the OneTouch Ultra Plus™ meter yielded accurate glucose values. For plasma glucose measurements, blood was collected by submandibular vein puncture using 5 mm Goldenrod® lancets into EDTA-coated microtubes. Samples were centrifuged at 2,000 × g for 10 minutes at 4°C, and the plasma fraction was applied to the glucose meter. These values are reported in the text as “plasma glucose,” while “blood glucose” refers to values obtained from whole-blood samples.

### Glucose tolerance test

Glucose tolerance tests (GTTs) were performed by intraperitoneal injection of a glucose bolus (2 g/kg body weight), followed by serial blood glucose measurements at baseline (prior to injection) and at 15, 30, 60, 90, and 120 minutes post-injection. Each mouse’s baseline value was used as its individual reference point. The area under the curve (AUC) for blood glucose over time, relative to the baseline, was calculated using GraphPad Prism software. Individual AUCs were grouped by experimental condition to assess the impact of interventions on glucose tolerance. A reduced mean AUC compared to control mice indicated improved glucose tolerance, while an increased AUC was interpreted as impaired glucose tolerance. To visualize changes in glucose tolerance over time, AUC values were expressed as fold changes relative to the control condition within each experiment.

### Insulin tolerance test

Insulin tolerance tests (ITTs) were performed by intraperitoneal injection of a human insulin bolus (1 IU/kg body weight), followed by serial blood glucose measurements at baseline (prior to injection) and at 15, 30, 60 and 90 minutes post-injection. Each mouse’s baseline value was used as its individual reference point. The area above the curve (referred as AUC for simplicity) for blood glucose over time, relative to the baseline, was calculated using GraphPad Prism software. Individual AUCs were grouped by experimental condition to assess the impact of interventions on insulin tolerance. A reduced mean AUC compared to control mice indicated impaired insulin tolerance, while an increased AUC was interpreted as improved insulin sensitivity. To visualize changes in insulin tolerance over time, AUC values were expressed as fold changes relative to the control condition within each experiment.

### Pyruvate tolerance test

Pyruvate tolerance tests (PTTs) were performed by intraperitoneal injection of a pyruvate bolus (2 g/kg body weight), followed by serial blood glucose measurements at baseline (prior to injection) and at 15-, 30-, 60-, 90-, and 120-minutes post-injection. Each mouse’s baseline value was used as its individual reference point. The area under the curve (AUC) for blood glucose over time, relative to the baseline, was calculated using GraphPad Prism software. Individual AUCs were grouped by experimental condition to evaluate the effect of interventions on pyruvate tolerance. A lower mean AUC compared to control mice was interpreted as a reduced gluconeogenic capacity (i.e., diminished glucose production from pyruvate), while a higher AUC indicated an enhanced gluconeogenic response.

### FDG PET-CT scan analysis

FDG PET-CT scans acquired for our previous study on hypoxic fuel rewiring were re-analyzed by Amide software to estimate the relative contribution of major organs to the overall increase in glucose uptake under hypoxia adaptation. Male mice were housed under normoxic (21% O_2_, *n* = 8) or hypoxic (8% O_2_, *n* = 5) conditions for 3 weeks. At the end of the exposure period, all animals received an intravenous injection of 2-deoxy-2-[¹⁸F]fluoro-D-glucose (FDG). FDG accumulation was used as a surrogate marker for tissue-specific glucose uptake.

Whole-body ¹⁸F signal was quantified by drawing same-size cylindrical regions of interest (ROIs) (“Whole-body ROI: 27, 27, 90 mm-elliptic cylinder) encompassing individual animals. To correct for urinary excretion of FDG, signal within the bladder was excluded from the total signal (ROI: 9, 9, 9 mm-ellipse). Organ-specific FDG uptake was determined by placing same-size elliptical ROIs over the heart (ROI: 10, 8, 10 mm-ellipse), brain (ROI: 9, 16, 10 mm-ellipse), and brown adipose tissue (BAT) (ROI: 9, 9, 9 mm-ellipse), guided by anatomical landmarks visible on the corresponding CT scans. The increase in whole-body glucose uptake under hypoxia was calculated as the difference in the percentage of injected dose (%ID) between hypoxic mice and the normoxia average (Δ%ID = %ID₈_%_ - %ID_2_₁_%_). The relative contribution of each organ to the overall Δ%ID was then computed to identify potential drivers of increased glucose uptake and represented in a pie chart.

### Hematological analysis

Complete blood counts were performed using a VetScan HM5 hematology analyzer (Abaxis, Union City, CA). Blood samples were collected via submandibular vein puncture using 5 mm Goldenrod® lancets and immediately transferred into EDTA-coated microtubes to prevent coagulation. Samples were stored at 4°C and analyzed within 1 to 6 hours of collection.

For analysis, 50 µL of EDTA-anticoagulated blood was automatically drawn by the VetScan HM5. The sample was split into two fractions. The first fraction was used for red blood cell (RBC) parameter analysis via impedance technology, providing measurements such as RBC count (10¹²/L) and mean corpuscular volume (MCV, fL). Hematocrit (%) was calculated as the product of RBC count and MCV divided by 10, following the formula: Hematocrit = (RBC × MCV) / 10. The second blood fraction was subjected to red blood cell lysis to enable quantification of non-RBC parameters. White blood cell (WBC) and platelet counts (10⁹/L) were determined by impedance in this lysed fraction. Hemoglobin concentration (g/dL) was measured photometrically at a wavelength of 540 nm using the same lysed sample.

### Phlebotomy Model

To achieve sustained red blood cell (RBC) reduction, serial phlebotomy was performed on mice housed under hypoxic (8% O_2_, n=8) and normoxic (21% O_2_, n=8) conditions. Every three days, 15% of the estimated total blood volume (TBV) was withdrawn via submandibular vein puncture using 5 mm Goldenrod® lancets. To maintain circulatory volume, an equal volume of sterile saline was administered intraperitoneally following each blood draw. Phlebotomized mice were compared to control groups of non-phlebotomized mice maintained under hypoxia (n=5) or normoxia (n=5).

Blood samples were collected on days 9, 15, and 21 of exposure for complete blood count (CBC) and plasma glucose analysis. Blood was collected into EDTA-coated microtubes and stored at 4°C until processing. Samples were split into two aliquots upon collection. One aliquot was used for plasma glucose measurement. Plasma was extracted by centrifugation within 1 hour of collection, and glucose levels were subsequently measured. The second aliquot was used for hematological analysis and processed within 1 to 6 hours of collection, as described in the CBC analysis section above.

### RBC Transfusion Model

To increase circulating RBC levels, mice housed under normoxic conditions (21% O_2_) were transfused with RBCs derived from donor mice previously exposed to either hypoxia (8% O_2_) or normoxia (21% O_2_) for four weeks. Recipient groups included mice receiving hypoxic RBCs (n=5), normoxic RBCs (n=5), or vehicle control (saline, n=5). Blood from donor mice was collected into tubes containing citrate buffer (pH 7.0) to a final concentration of 0.3% to prevent coagulation. Samples were thoroughly mixed and centrifuged at 2,000 × g for 10 minutes at 4°C. The plasma and buffy coat layers were discarded to isolate a clean pellet of packed RBCs. This pellet was then pooled and resuspended in sterile saline to achieve a 75% hematocrit (i.e., 75% packed RBCs by volume).

Recipient mice received retro-orbital injections containing 2000 µL of 75%-packed RBCs twice daily for two consecutive days. Endpoint blood analysis was conducted the following day. Blood was collected via submandibular vein puncture into EDTA-coated microtubes and analyzed immediately for hematological parameters and plasma glucose as described above.

### Type I Diabetes Model

Type I diabetes was induced in 8-week-old male mice by intraperitoneal (i.p.) injection of streptozotocin (STZ; n=16) or vehicle (citrate buffer, pH 4.5; n=16) for five consecutive days. Two weeks post-injection, blood glucose levels and body weight were assessed to confirm the onset of hyperglycemia. Following verification of diabetes induction, STZ-treated hyperglycemic mice and vehicle-treated controls were randomly assigned to hypoxia or normoxia groups. Mice were exposed to either hypoxic (8% O_2_; n=8 STZ-treated, n=8 vehicle-treated) or normoxic (21% O_2_; n=8 STZ-treated, n=8 vehicle-treated) conditions for a period of three weeks. Throughout the hypoxia/normoxia exposure, blood glucose levels and body weight were monitored every three days to evaluate the physiological effects of hypoxia in diabetic and non-diabetic mice.

### HFD-Induced Hyperglycemia Model

Male C57BL/6J mice fed either a high-fat diet (HFD; JAX® Diet-Induced Obese, stock #380050, n=11) or standard chow diet (CD; stock #000664, n=16) were purchased from The Jackson Laboratory. Upon arrival, mice were allocated to receive either daily oral gavage with HypoxyStat (600 mg/kg) or vehicle for a duration of 2.5 weeks. Group allocation was as follows: HFD-fed mice received either HypoxyStat (n=5) or vehicle (n=6), while CD-fed mice received either HypoxyStat (n=8) or vehicle (n=8). Blood glucose levels were measured after two weeks of daily dosing. Glucose tolerance was evaluated after two and a half weeks of treatment to assess the metabolic effects of HypoxyStat in both dietary contexts.

### *In vivo* 2-deoxy-D-glucose (U-^13^C) uptake assay

2-deoxy-D-glucose (2DG) (U-^13^C) (Cambridge Isotope Laboratories) was freshly dissolved in sterile water on the day of the experiment. Mice exposed to hypoxia (8% O_2_, n=5) or normoxia (21% O_2_, n=5) for three weeks were administered a bolus injection of 2DG (U-^13^C) at a dose of 1 g/kg body weight via the retro-orbital route. Blood samples were collected at 2-, 10-, 30-, and 120-minutes post-injection in EDTA-coated microtubes. Collected EDTA-blood was stored at 4°C and processed at the end of the 2-hour experimental period. Blood samples collected at 2 minutes were split into two aliquots upon collection. One aliquot was used for hematological analysis, to ensure proper normalization of 2DG uptake by the total number of RBC per sample, using the formula: Volume of analyzed RBC (20 μL) × 10^9^ / MCV (fl) = Number of analyzed RBCs in 20 μL

The second aliquot was processed together with the rest of the samples. Blood samples were centrifuged at 2,000 × *g* for 10 minutes at 4°C to separate plasma and cellular components. A 5 μL aliquot of plasma was transferred to a clean microtube and stored at −80°C for later analysis. The remaining plasma and buffy coat were discarded to isolate a clean pellet of packed red blood cells (RBCs). A 5 μL RBC pellet was collected and stored at −80°C for subsequent isotopic analysis.

### *Ex vivo* glucose uptake assay

Blood samples were collected from mice exposed to hypoxia (8% O_2_, n=5) or normoxia (21% O_2_, n=5) for four weeks in EDTA-coated microtubes and stored on ice until processing. Blood samples were split into two aliquots upon collection. One aliquot was used for hematological analysis, to ensure proper normalization of glucose uptake by the total number of RBC per sample, using the formula: Volume of analyzed RBC (10 μL) × 10^9^ / MCV (fl) = Number of analyzed RBCs in 10 μL

The second aliquot was centrifuged at 2,000 g for 10 minutes at 4°C and all the plasma from normoxic mice was pooled in a vial. Plasma glucose in the pooled normoxic plasma was measured for reference using the OneTouch Ultra Plus^TM^ glucose meter. Upon plasma and buffy coat layers removal, 10 μL of RBCs pellet from each hypoxic and normoxic sample were incubated with 10 μL of the pooled normoxic plasma. These samples were incubated at 37°C for 10 minutes in a rocker. One sample only containing plasma was included to test non-RBC specific changes in glucose levels upon incubation. After this time, samples were centrifuged to isolate the plasma that had been in contact with the experimental RBCs. Remaining plasma glucose was measured in all samples using the OneTouch Ultra Plus^TM^ glucose meter and the value was compared to the plasma glucose level in the incubated sample without RBCs (234 mg/dL), which did not present a significant change in glucose levels during the incubation.

### *In vivo* glucose (U-^13^C) tracer experiment

Glucose (U-^13^C) (Cambridge Isotope Laboratories) was freshly dissolved in sterile water on the day of the experiment. Mice exposed to hypoxia (8% O_2_, n=5) or normoxia (21% O_2_, n=5) for three weeks were administered a bolus injection of glucose (U-^13^C) at a dose of 1 g/kg body weight via the retro-orbital route. Blood samples were collected at 2, 10, 30, and 120 minutes post-injection in EDTA-coated microtubes. Collected EDTA-blood was stored at 4°C and processed at the end of the 2-hour experimental period. Samples were centrifuged at 2,000 × *g* for 10 minutes at 4°C to separate plasma and cellular components. A 5 μL aliquot of plasma was transferred to a clean microtube and stored at −80°C for later analysis. The remaining plasma and buffy coat were discarded to isolate a clean pellet of packed red blood cells (RBCs). A 5 μL RBC pellet was collected and stored at −80°C for subsequent isotopic analysis.

### Polar metabolites extraction

Plasma was extracted from blood by centrifuging at 2,000 g for 10 minutes at 4°C and collecting supernatant. For tracer experiments, plasma metabolites were extracted by adding 45 μL of 80% methanol to every 5 μL of plasma. Samples were vortexed for 10s and incubated at −80°C for 20 minutes. Next, samples were centrifuged at 20,000g for 10 minutes at 4°C and supernatant was collected. Supernatants were subsequently lyophilized at 4°C and resuspended in 40 μL 60% ACN. Samples were sonicated, mixed in a thermomixer and incubated for 20 minutes at 4°C. Samples were centrifuged at 20,000 g for 20 minutes at 4°C and supernatants were transferred to suitable vials for LC-MS injection.

For RBC polar metabolites extraction, 400 μL of 40/40/20 solvent was added to 20 μL of isolated RBCs. Samples were mixed in a thermomixer, sonicated and incubated for 1 minute in liquid nitrogen twice. After the second cycle, samples were incubated in ice for 20 minutes and centrifuged at 20,000 g for 20 minutes at 4°C. Supernatants were subsequently lyophilized at 4°C and resuspended in 100 μL 60% ACN. Samples were sonicated, mixed in a thermomixer and incubated for 20 minutes at 4°C. Samples were centrifuged at 20,000 g for 20 minutes at 4°C and supernatants were transferred to suitable vials for LC-MS injection.

### LC-MS

Samples were run on an Orbitrap Exploris 240 (OE240) high resolution mass spectrometer (HRMS, Thermo Fisher Scientific) using electrospray ionization. The OE240 was coupled to hydrophilic interaction chromatography on a Vanquish Horizon ultra-high performance liquid chromatography (UHPLC, Thermo Fisher Scientific). 2 μL of polar metabolite samples were injected into the LC-MS system and separated on a iHILIC-(P) Classic column (2.1×150 mm, 5 µm; part # 160.152.0520; HILICON AB). The autosampler was maintained at 4°C and the column was maintained at 40°C during runs. Mobile phase A was 20 mM ammonium bicarbonate in water, with ammonium hydroxide added to reach a pH of 9.6. Mobile phase B was acetonitrile. Flow rate was 200 μL/min. The gradient was the following: 0-20 min: 80% B to 20% B; 20-20.5 min: linear gradient from 20-80% B; 20.5-28 min: stable at 80% B.77 A 10-minute equilibration phase was included at starting conditions before each injection. The OE240 ran in full-scan, polarity-switching mode with an ion voltage of 3.5 kV in positive mode and 3.25 kV in negative mode, and the scan range was 70-1000 m/z. The orbitrap resolution was 60000, RF Lens was 60%, AGC target was 1e7, and the maximum injection time was 200 ms. The sheath gas was set to 35 units, auxiliary gas was 10 units, and sweep gas was 0.5 units. The ion transfer tube temperature was 300°C, and the vaporizer temperature was 35°C.

### Statistical analysis

Sample sizes for each measurement were based on previously reported results and varied depending on the data type and study design. Data are presented as mean ± standard deviation (SD) in dot plots unless otherwise specified. For comparisons between two groups, either paired or unpaired *t*-tests were used depending on the experimental design. When data met assumptions of normality, a *t*-test was applied; otherwise, a non-parametric Mann–Whitney test was used. For comparisons involving more than two groups, one-way analysis of variance (ANOVA) followed by Dunnett’s multiple comparisons test was performed. When data involved two categorical independent variables (e.g., time and genotype, or genotype and treatment), a two-way ANOVA or a mixed-effects model (with Geisser–Greenhouse correction) was applied. Specifically, mixed-effects models were used when repeated measurements were taken from the same group of individuals over time, while two-way ANOVA was used when each measurement came from distinct individuals. Statistical outliers were identified using Grubb’s test (extreme studentized deviate method). All statistical analyses were performed using Prism 10 (GraphPad Software, California, USA), unless otherwise noted. A *p*-value < 0.05 was considered statistically significant.

## Figures

Figures were generated using Prism 10 (GraphPad Software, California, USA) and Adobe Illustrator. Biorender was used to create new diagrams.

## RESOURCE AVAILABILITY

### LEAD CONTACT

Additional information or requests for resources will be fulfilled by the lead contact, Isha Jain.

### MATERIAL AVAILABILITY

No new materials were created for this study.

### DATA AND CODE AVAILABILITY

No new code was generated for this study. No large datasets were generated for this study.

## ACKNOWLEDGEMENTS

We appreciate thoughtful discussions with members of the Jain Lab including Jonathan Tai, Alan Baik and Brandon Desousa. We are deeply grateful to Jessica Beserra Felix, Brandon Chew and Tej Joshi for their invaluable technical assistance and support. We gratefully acknowledge the support of the Yamanaka Lab for providing access to the blood analyzer and for their generous technical assistance. We are deeply grateful to Brian Plosky for thoughtful editorial input and valuable feedback on the manuscript. We sincerely thank Chiara Ricci-Tam for the design and refinement of figures and the graphical abstract. IHJ was supported by NIH DP5 DP5OD026398. YMM was supported by the CIRM postdoctoral fellowship. This work was supported by NIH DP5OD026398, a gift from Dave Wentz and the Hillblom Foundation Award.

## AUTHOR CONTRIBUTIONS

IHJ and YMM conceived the project. All authors designed, performed and analyzed experiments. IHJ and YMM wrote and edited the manuscript with final approval from all co-authors

## DECLARATION OF INTERESTS

IHJ has patents related to hypoxia therapy.

## Supplementary Figures

**Figure S1.**
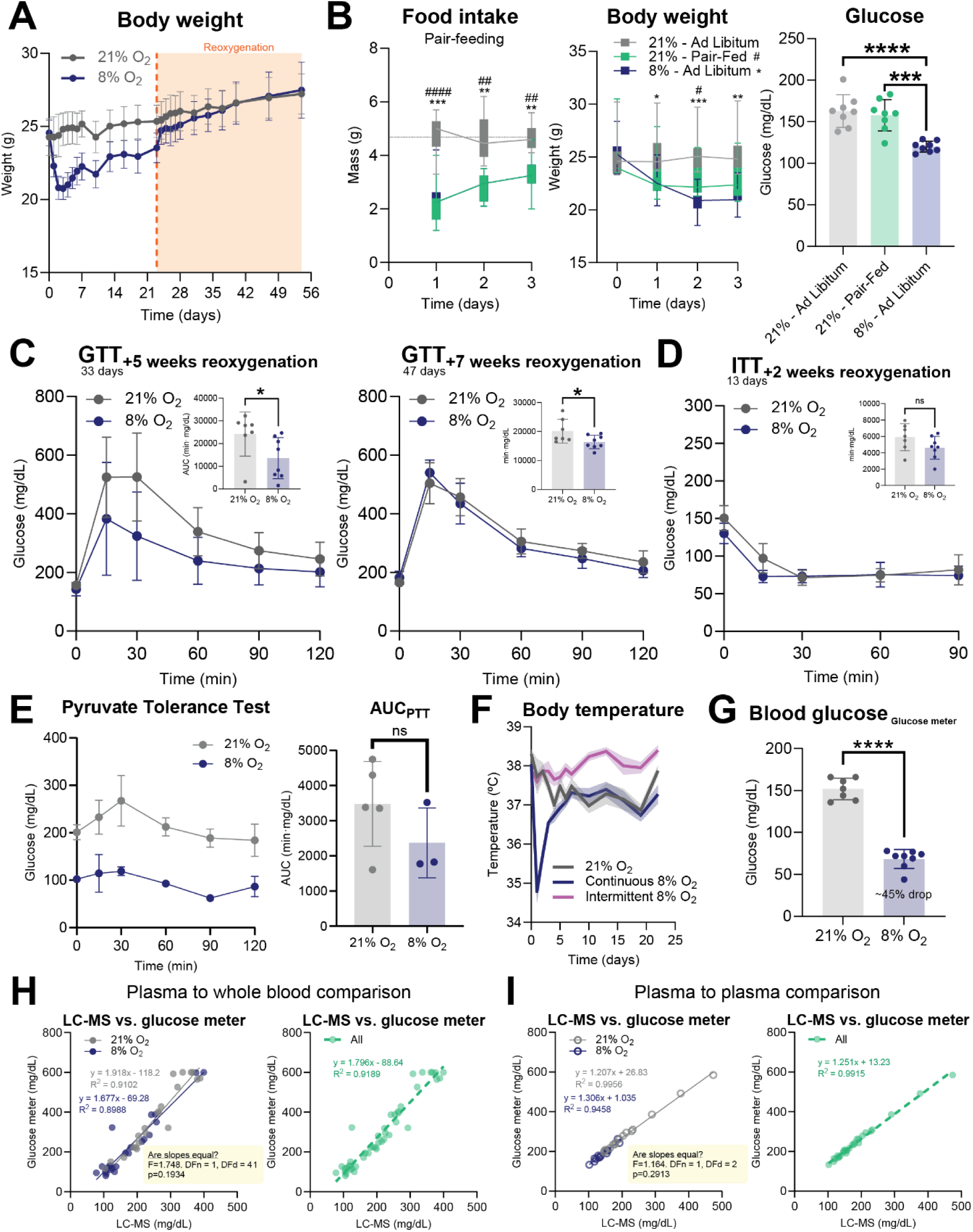
Hypoxia leads to improved glucose tolerance, which is not fully explained by increased glucose uptake by tissues. **(A)** Body weight over time in hypoxia (8% O_2_, n=8) or normoxia (21% O_2_, n=7) and upon subsequent reoxygenation, marked with an orange dashed line at day 23. Mean ± SD are shown. **(B)** Food intake per mouse and body weight over time with hypoxic pair-feeding in normoxia (21% O_2_ Pair-Fed, n=8) compared to *ad libitum* feeding in hypoxia (8% O_2_ Ad Libitum, n=8) and normoxia (8% O_2_ Ad Libitum, n=8) (graphs on the left). Dunnett’s multiple comparisons test was used. Endpoint blood glucose in pair-fed mice vs. hypoxic and normoxic ad libitum-fed mice (right). Ordinary one-way ANOVA was used. **(C)** Glucose tolerance tests after five weeks (33 days, left) and seven weeks (47 days, right) of reoxygenation in mice that had previously been exposed to hypoxia (8% O_2_, n=8) for three weeks or normoxia (21% O_2_, n=7). Mean ± SD are shown. AUC values from individual curves were plotted for each test (top right of both graphs) and analyzed by an unpaired two-tailed Student’s t-test. **(D)** Insulin tolerance test after two weeks (13 days) of reoxygenation in mice that had previously been exposed to hypoxia (8% O_2_, n=8) for three weeks or normoxia (21% O_2_, n=7). Mean ± SD are shown. AUC values from individual curves were plotted (top right of both graphs) and analyzed by an unpaired two-tailed Student’s t-test. **(E)** Pyruvate tolerance test after two weeks in hypoxia (8% O_2_, n=3) or normoxia (21% O_2_, n=5) (left) and AUC values from individual curves were plotted and analyzed by an unpaired two-tailed Student’s t-test (right). **(F)** Rectal temperature over time in intermittent hypoxia (8% O_2_ for 8 hours and 21% O_2_ for 16 hours, n=8), continuous hypoxia (continuous 8% O_2_, n=8) or normoxia (continuous 21% O_2_, n=8). Mean ± SD are shown. **(G)** Tail vein blood glucose values measured by glucose meter from mice submitted to three weeks of hypoxia (8% O_2_, n=8) or normoxia (21% O_2_, n=7). Mean ± SD are shown. Unpaired two-tailed Student’s t-test was used. **(H)** Simple linear regression of glucose values obtained by LC-MS (on the plasma fraction) or by a glucose meter (on the whole-blood fraction) from the same blood samples coming from mice exposed to hypoxia for one or three weeks (8% O_2_, n=12 and n=12) or normoxia (21% O_2_, n=12 and n=9) considering their oxygen tension differences (left) or plotting all the values together (right). **(I)** Simple linear regression of glucose values obtained by LC-MS (on the plasma fraction) or by a glucose meter (also on the plasma fraction) from the same blood samples coming from mice exposed to hypoxia for three weeks (8% O_2_, n=16) or normoxia (21% O_2_, n=16) considering their oxygen tension differences (left) or plotting all the values together (right). ∗p < 0.05, ∗∗p < 0.01, ∗∗∗p < 0.001, and ∗∗∗∗p < 0.0001.

**Figure S2.**
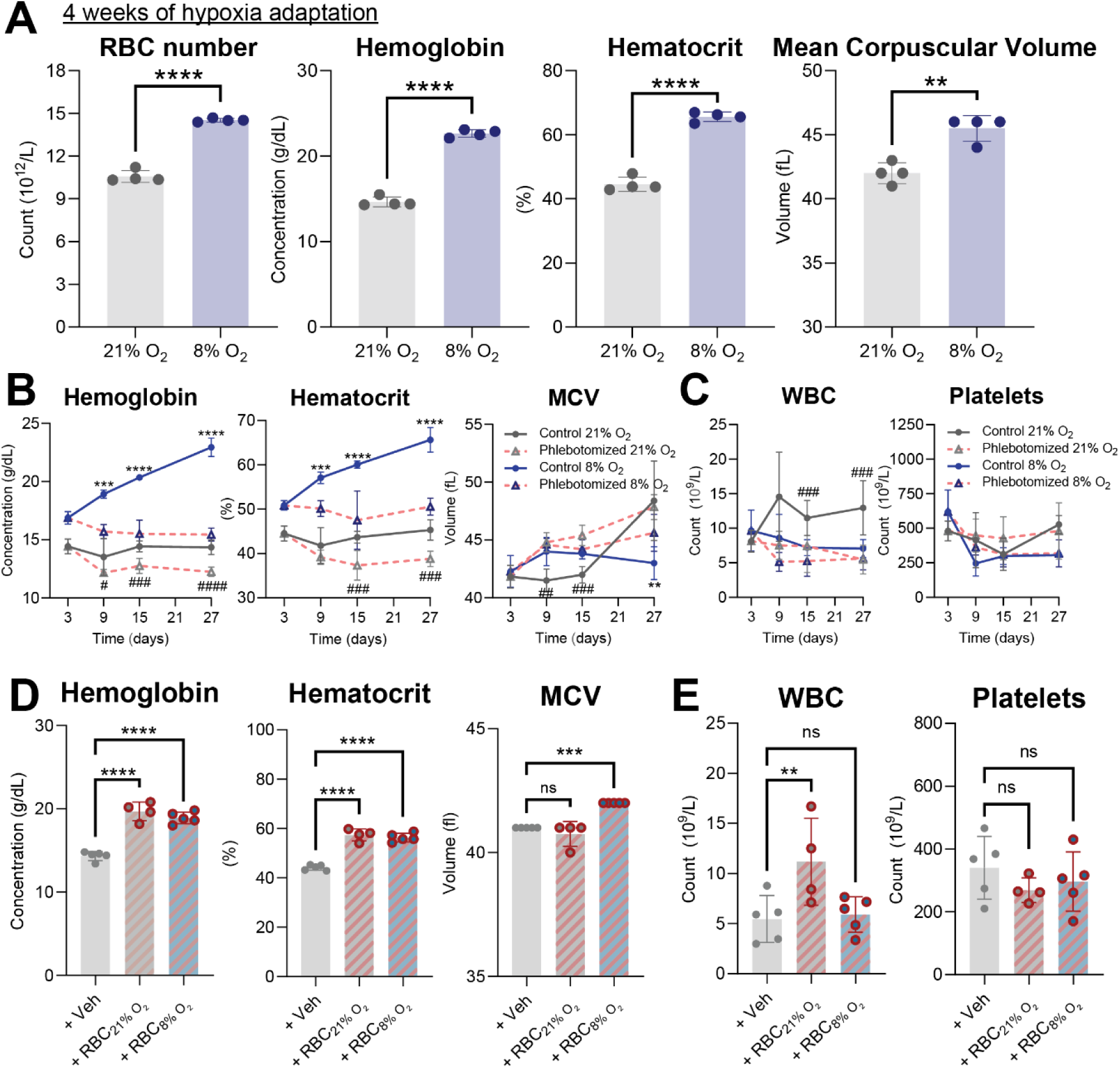
Erythrocytosis is necessary and sufficient to explain hypoglycemia in hypoxia. **(A)** RBC count, hemoglobin concentration, hematocrit and mean corpuscular volume (MCV) values in mice after four weeks of hypoxia (8% O_2_, n=4) or normoxia (8% O_2_, n=4), as evidence for erythrocytosis during chronic hypoxic adaptation. **(B)** Hemoglobin concentration, hematocrit and MCV over time in hypoxia (8% O_2_ control and phlebotomized, n=5 and n=8, respectively) or normoxia (21% O_2_ control and phlebotomized, n=5 and n=8, respectively). Mean ± SD are shown. Two-way ANOVA was used. **(C)** WBC and platelets counts over time in hypoxia (8% O_2_ control and phlebotomized, n=5 and n=8, respectively) or normoxia (21% O_2_ control and phlebotomized, n=5 and n=8, respectively). Mean ± SD are shown. Two-way ANOVA was used. **(D)** Endpoint hemoglobin, hematocrit and MCV after RBC transfusion in recipient mice injected with RBCs coming from hypoxic (8% O_2_, to n=5 recipients) or normoxic animals (21% O_2_, to n=5 recipients) as well as injected with vehicle (saline, to n=5 recipients). Ordinary one-way ANOVA was used. **(E)** Endpoint WBCs and platelets counts after RBC transfusion in recipient mice injected with RBCs coming from hypoxic (8% O_2_, to n=5 recipients) or normoxic animals (21% O_2_, to n=5 recipients) as well as injected with vehicle (saline, to n=5 recipients). Ordinary one-way ANOVA was used. ∗p < 0.05, ∗∗p < 0.01, ∗∗∗p < 0.001, and ∗∗∗∗p < 0.0001. (Phlebotomized to control comparisons at 8% O_2_). #p < 0.05, ##p < 0.01, ###p < 0.001, and ####p < 0.0001. (Phlebotomized to control comparisons at 21% O_2_)

**Figure S3.**
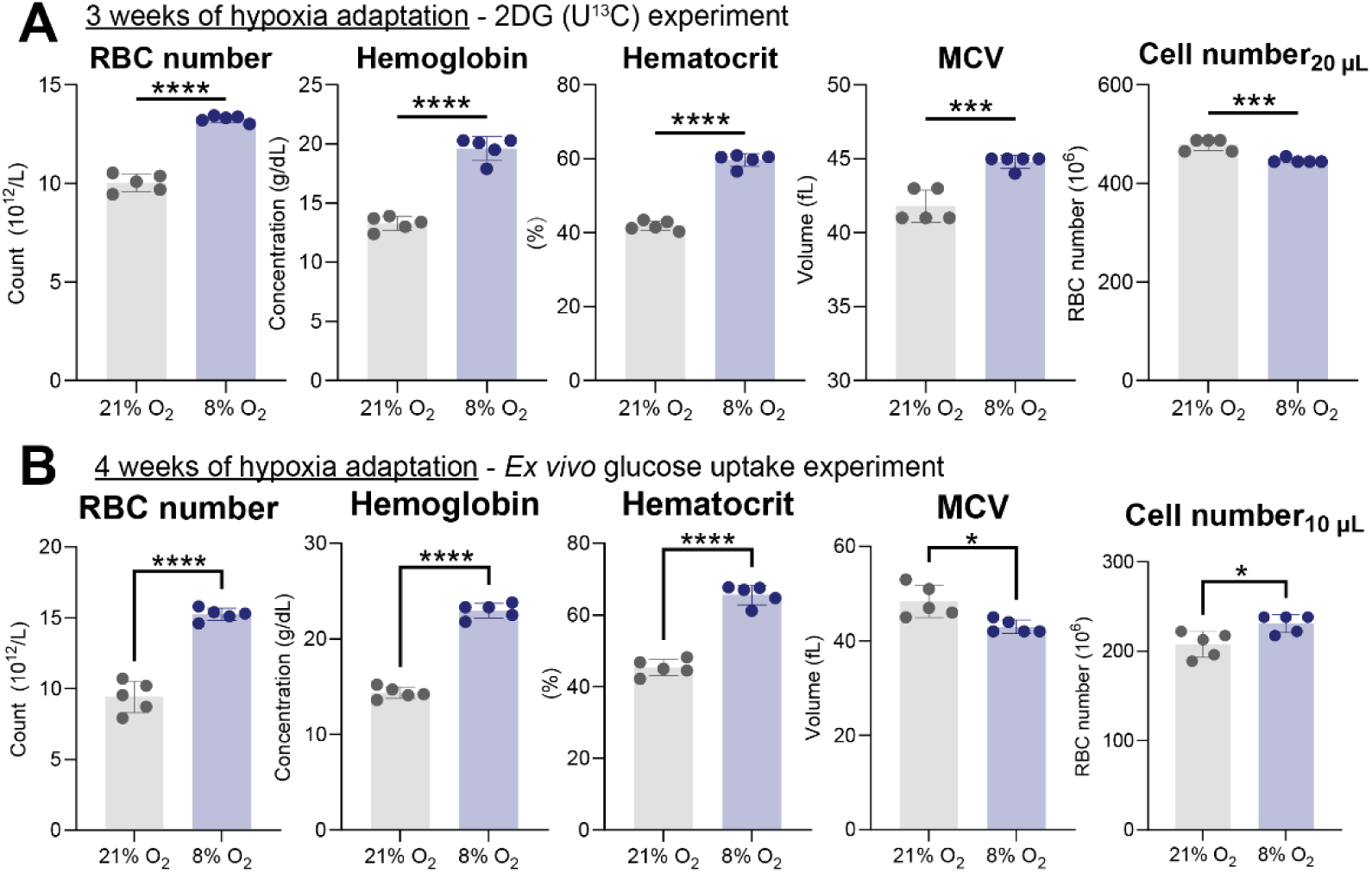
Glucose uptake by RBCs multiplies 3-fold in hypoxia to build hemoglobin allosteric regulator 2,3-DPG. **(A)** RBC count, hemoglobin concentration, hematocrit, MCV and estimated number of RBCs in the preparations for LC-MS quantification from mice exposed to hypoxia (8% O_2_, n=5) or normoxia (21% O_2_, n=5) for three weeks and used for the *in vivo* 2DG-based glucose uptake experiment. Unpaired two-tailed Student’s t-test was used. **(B)** RBC count, hemoglobin concentration, hematocrit, MCV and estimated number of RBCs in the preparations for the *ex vivo* glucose uptake experiment from mice exposed to hypoxia (8% O_2_, n=5) or normoxia (21% O_2_, n=5) for three weeks. Unpaired two-tailed Student’s t-test was used. ∗p < 0.05, ∗∗p < 0.01, ∗∗∗p < 0.001, and ∗∗∗∗p < 0.0001.

**Figure S4.**
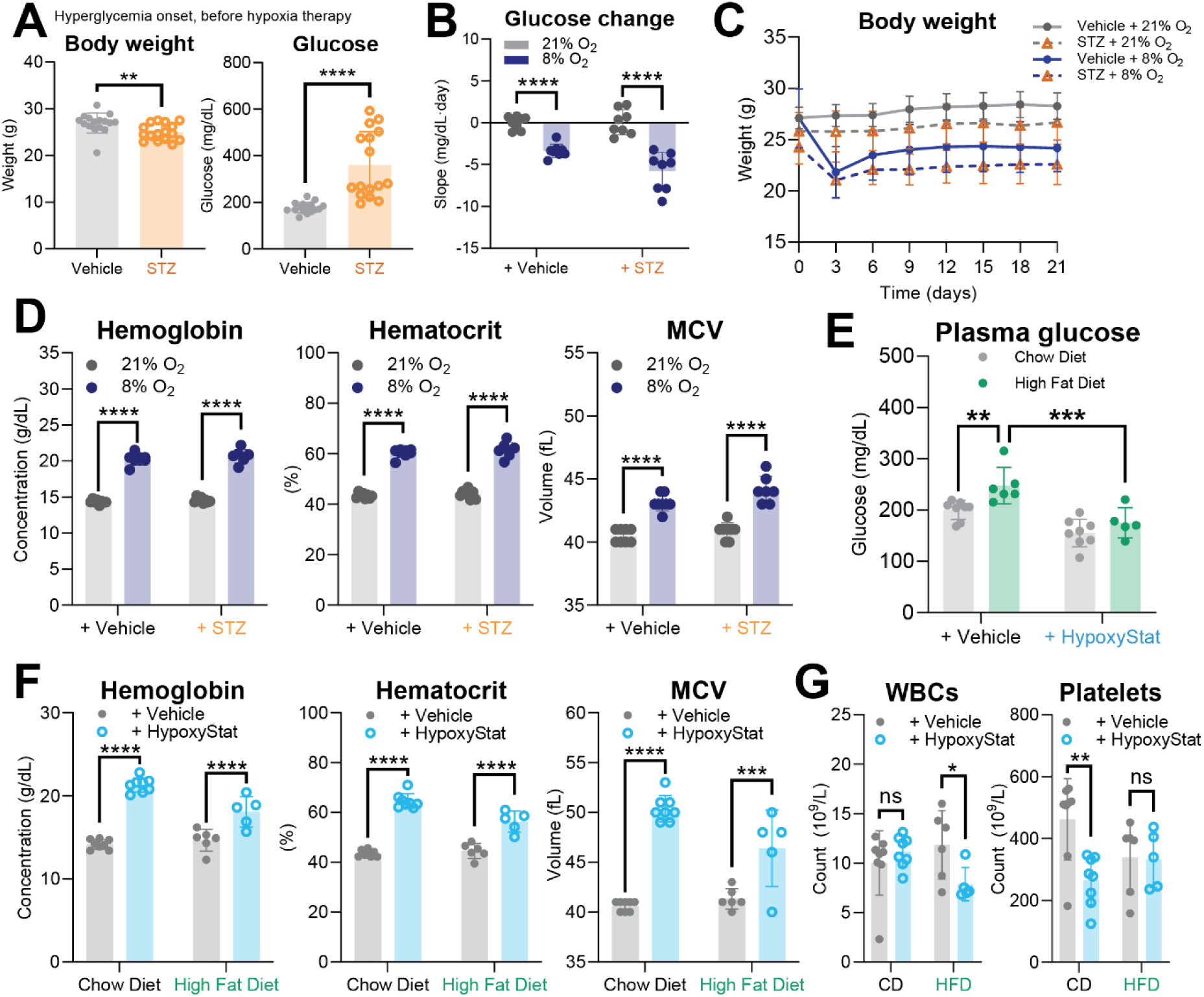
Hypoxia ameliorates STZ- and HFD-induced hyperglycemia. **(A)** Body weight (left) and blood glucose (right) two weeks after STZ-treatment. Unpaired two-tailed Student’s t-test was used. **(B)** Slopes from the linear regression models behind the blood glucose drop in hypoxic mice over time, including vehicle- and STZ-treated animals. Unpaired two-tailed Student’s t-test was used. **(C)** Body weight over time in hypoxia (8% O_2_ Vehicle- or STZ-treated mice, n=8 and n=8, respectively) or normoxia (21% O_2_ Vehicle- or STZ-treated mice, n=8 and n=8, respectively). Mean ± SD are shown. **(D)** Endpoint hemoglobin, hematocrit and MCV values after three weeks in hypoxia (8% O_2_ Vehicle- or STZ-treated mice, n=8 and n=7, respectively) or normoxia (21% O_2_ Vehicle- or STZ-treated mice, n=8 and n=8, respectively). Sidak’s multiple comparison test was used. **(E)** Endpoint plasma glucose in HypoxyStat-treated mice (HypoxyStat CD- or HFD-fed, n=8 and n=5, respectively) or vehicle-treated animals (Vehicle CD- or HFD-fed, n=8 and n=6, respectively). Sidak’s multiple comparison test was used. **(F)** Endpoint hemoglobin, hematocrit and MCV values after HypoxyStat dosing (HypoxyStat with CD or HFD, n=8 and n=5, respectively) or vehicle-dosing (Vehicle with CD or HFD, n=8 and n=6, respectively). Sidak’s multiple comparison test was used. ∗p < 0.05, ∗∗p < 0.01, ∗∗∗p < 0.001, and ∗∗∗∗p < 0.0001.

